# RAD50 promotes DNA repair by homologous recombination and restrains antigenic variation in African trypanosomes

**DOI:** 10.1101/2020.03.17.994905

**Authors:** Ann-Kathrin Mehnert, Marco Prorocic, Annick Dujeancourt-Henry, Sebastian Hutchinson, Richard McCulloch, Lucy Glover

## Abstract

Homologous recombination dominates as the major form of DNA repair in *Trypanosoma brucei*, and is especially important for recombination of the subtelomeric variant surface glycoprotein during antigenic variation. RAD50, a component of the MRN complex (MRE11, RAD50, NBS1), is central to homologous recombination through facilitating resection and governing the DNA damage response. The function of RAD50 in trypanosomes is untested. Here we report that RAD50 is required for RAD51-dependent homologous recombination, phosphorylation of histone H2A and controlled resection following a DNA double strand break (DSB). Perhaps surprisingly, DSB resection in the *rad50* nulls was not impaired and appeared to peak earlier than in the parental strains. Finally, we show that RAD50 suppresses DNA repair using donors with short stretches of homology at a subtelomeric locus, with null strains producing a greater diversity of expressed VSG variants following DSB repair. We conclude that RAD50 promotes stringent homologous recombination at subtelomeric loci and restrains antigenic variation.

## INTRODUCTION

*Trypanosoma brucei* (*T. brucei*) is a protozoan parasite and the causative agent of human African trypanosomiasis, or sleeping sickness, and nagana in cattle. Trypanosomes cycle between their insect vector, the tsetse fly, and mammalian hosts, where they colonise the blood, fat^1^ and skin^2^ and eventually cross the blood brain barrier in late stage infection. If left untreated, trypanosomiasis is normally fatal^3^. In the mammalian host, each trypanosome cell is covered in a dense layer of a single species of variant surface glycoprotein (VSG). The highly immunogenic VSG layer^4,5^ acts as an barrier, concealing other surface components from the host immune response ^6^. Trypanosomes maintain a persistent infection by continuously escaping the host’s immune response though antigenic variation^7^. Central to this survival strategy is monoallelic expression of the VSG from a subtelomeric locus, known as an expression site (*VSG*-ES), and stochastic VSG switching. The ~ 15 *VSG*-ESs in the trypanosome genome share a high degree of sequence and structure conservation ^8^, each being an RNA polymerase-I (RNA Pol-I) polycistronic transcription unit with a single *VSG* gene found adjacent to the telomere, up to 60 kb downstream of the promoter ^8^. The *VSG* gene is flanked by two sets of repetitive sequence:downstream is the telomere, and upstream is a block of repetitive sequence, known as the 70-bp repeats, which modulates *VSG* switching ^8,9^. Characteristic of a trypanosome infection are recrudescent waves of parasitemia, each of which is composed of a diverse *VSG* expressing population, with between 7 - 79 VSGs detected in each peak of parasitemia ^10–12^. VSG diversity arises through altering the single *VSG*-ES that is transcribed or, more commonly, by recombination of silent *VSGs* into the active *VSG*-ES. The seemingly unrestricted use of *VSG* genes might be expected to result in a rapid exhaustion of the *VSG* gene repertoire. However, the parasite’s ability to sustain an infection appears to lie in an enormous repertoire of >2000 *VSG* genes and pseudogenes^13–15^, mainly found in subtelomeric *VSG* arrays, and a capacity for generation of novel ‘mosaic’ *VSG* genes through segmental gene conversion of multiple (pseudo) VSGs, in particular late in infection ^10,11,14^. Importantly, almost all of the array *VSGs* are associated with upstream tracts of 70-bp repeats, providing the necessary substrate needed for homologous recombination mediated antigenic variation ^16^.

A DNA double-strand break (DSB) is an extremely toxic lesion in any cell, which if left unrepaired can lead to cell death. In *T. brucei* RAD51-dependent homologous recombination (HR) dominates as the major DNA repair and recombination pathway, with microhomology mediated end-joining (MMEJ) playing a minor role^17–20^. HR is important for *VSG* switching, and though it is not clear how MMEJ acts in this reaction, repair of induced DSBs can occur by coupled HR and MMEJ, and MMEJ is more frequently used for repair of DSBs induced within the *VSG*-ES^21^. Unrepaired DSBs appear to persist throughout the cell cycle without inhibiting the trypanosomes ability to replicate their DNA^22^, but whether HR or MMEJ are regulated is unknown. In addition, non-homologous end-joining (NHEJ) appears to be absent in trypanosomes^21,23,24^. These features of trypanosome DSB repair contrast with mammalian cells, where NHEJ is highly active, HR predominates in S and G_2_ phase cells and MMEJ is considered a minor reaction^25^. In trypanosomes both transcriptionally active and silent subtelomeres are fragile ^26,27^, and accumulate natural breaks. Within the active *VSG*-ES specifically, a DSB between the *VSG* and 70-bp repeats acts as a potent driver of antigenic variation and precipitates VSG switching^27^. Several DNA repair and recombination proteins have been shown to be important for antigenic variation in trypanosomes, thus linking VSG switching with this process:loss of RAD51^18^, the RAD51-3 paralogue^28^, or the RAD51-interacting protein BRCA2 ^29,30^ results in impaired VSG switching, while loss of RECQ2^31^, TOPO3α or RMI1 increases VSG switching^32,33^, as does loss of the histone variants H3.V and H4.V^15^. Loss of ATR, which is involved in DNA damage signalling, impairs monoallelic *VSG* expression and increases VSG switching through localized DNA damage^34^. Histone Acetyltransferase (HAT3) is required for recombination repair of a chromosome-internal DSB, but suppresses DSB repair within the *VSG*-ES which suggests repair is compartmentalised in trypanosomes^35^.

The DNA damage response (DDR) is an orchestrated cellular response to many different genome lesions, including DSBs, which most commonly form via stalled replication forks ^36^. Critical to DSB repair is the MRE11 - RAD50 - NBS1 (MRN) complex (in yeast MRE11 – RAD50 – XRS1, MRX), which acts as a DNA damage sensing complex and is responsible for recognizing the free DNA ends, where it is one of the first complexes to bind and initiate HR ^37,38^. MRE11 – RAD50 forms the core of this complex and is conserved across all domains of life, whereas NBS1 only forms part of the complex in eukaryotes ^37^. MRN consists of two molecules of each component protein, and diffuses along homoduplex DNA searching for free DNA ends – a process that is driven by RAD50 ^39^. The MRE11 subunit is a nuclease with both 5’ flap endonuclease activity and 3’⟶ 5’ exonuclease activity and catalyses resection through cleaving the 5’ strand, internal to the DSB, which is then resected using its exonuclease function to generate the short 3’ single-strand (ss) DNA overhangs ^40^. These overhangs are further resected by Exonuclease 1 (EXO1), forming long tracts of 3 ssDNA on either side of the DSB ^39^. NBS1, the eukaryote specific component, is responsible for binding multiple phosphorylated proteins and recruiting MRE11 and RAD50 to DSB sites ^41^ through its interaction with MRE11, CtIP, which is also required for initiating resection, and the ATM kinase ^42^. End recognition and DSB processing by MRN is an ATP dependent process:here, ATP binding to RAD50 acts to switch the complex from an open to a closed conformation ^43^, which facilitates DSB recognition, tethering and ATM activation ^43^. In yeast the MRX complex also acts in telomere maintenance by binding the end of short telomeres and recruiting TEL1, which then recruits telomerase to extend the telomere ^44^. Conversely, in mammalian cells, MRN regulates an ATM dependent response at dysfunctional telomeres ^45^.

RAD50 (Tb.927.11.8210), MRE11 (Tb927.2.4390) ^46,47^ and NBS1 (Tb 927.8.5710) homologues are present in the trypanosome genome and previous studies have shown that MRE11 is required for HR but its inactivation did not lead to telomere shortening or changes in VSG switching ^46,47^, despite the dominance of HR in repair in trypanosomes and requirement for the reaction in antigenic variation. What roles these proteins play in the trypanosome DDR is largely unexplored. In addition, though we know that DSBs accumulate at the subtelomeres ^26,27^, it is unclear how they are sensed or how they contribute to antigenic variation. Given the central, early role of the MRN complex in DSB recognition and in telomere maintenance we set out to characterise its role in HR and VSG switching in trypanosomes. We found that RAD50, like MRE11, is required for efficient HR, and in its absence MMEJ dominated as the major form of repair. RAD50 also plays a perhaps surprising role in VSG switching, where it restricts HR substrate selection in the *VSG* repertoire and so may act to preserve the *VSG* archive during long-term infections.

## MATERIALS AND METHODS

### *Trypanosoma brucei* growth and manipulation

Lister 427, MITat1.2 (clone 221a), bloodstream stage cells were cultured in HMI-11 medium ^79^ at 37.4 °C with 5 % CO_2._ Cell density was determined using a haemocytometer. For transformation, 2.5 × 10^7^ cells were spun for 10 minutes at 1000g at room temperature and the supernatant discarded. The cell pellet was resuspend in prewarmed cytomix solution ^80^ with 10 μg linearised DNA and place in a 0.2 cm gap cuvette, and nucleofected (Lonza) using the X-001 program. The transfected cells were placed into one 25 cm^2^ culture flask per transfection with 36 ml warmed HMI-11 medium only and place in an incubator to allow the cells to recover for approximately 6 hours. After 6 hours, the media distributed into 48-well plates with the appropriate drug selection. Strains expressing TetR and I-*Sce*I with I-*Sce*I recognition-sites at a chromosome-internal locus ^17^ and an active *VSG*-ESs ^27^ have been described previously. G418, and blasticidin were selected at 10 μg.ml^−1^ and 2 μg.ml^−1^ respectively. Puromycin, phleomycin, G418, hygromycin and blasticidin and tetracycline were maintained at 1 μg.ml^−1^. Clonogenic assay were plated out at either 32 cells per plate under both inducing and non-inducting conditions for ^1^HR and VSG^up^ strains and 480 cells per plate for VSG^up^ strains under inducing conditions. Plates were counted 5-6 days later and subclones selected for further analysis.

To generate the *RAD50* nulls in the ^1^HR strain we employed multi-step transfection strategy ^81^ that recycled a *Neomycin phosphotransferase* gene (*NEO*) in order to rescue one marker (here *Blasticidin - BLA*). Briefly, an I-*Sce*I recognition sites was inserted into the pRAD50-BLA knock-out cassette between the 5’UTR and *BLA* ORF (Figure 1C) in the 2T1 cell line ^82^ with a tetracycline inducible *Sce* ORF. Induction of Sce induces a break in the *BLA* cassette and subsequent repair, using homology in the *NEO* modified allele, replaces *BLA* with *NEO*.

**Figure 1:**
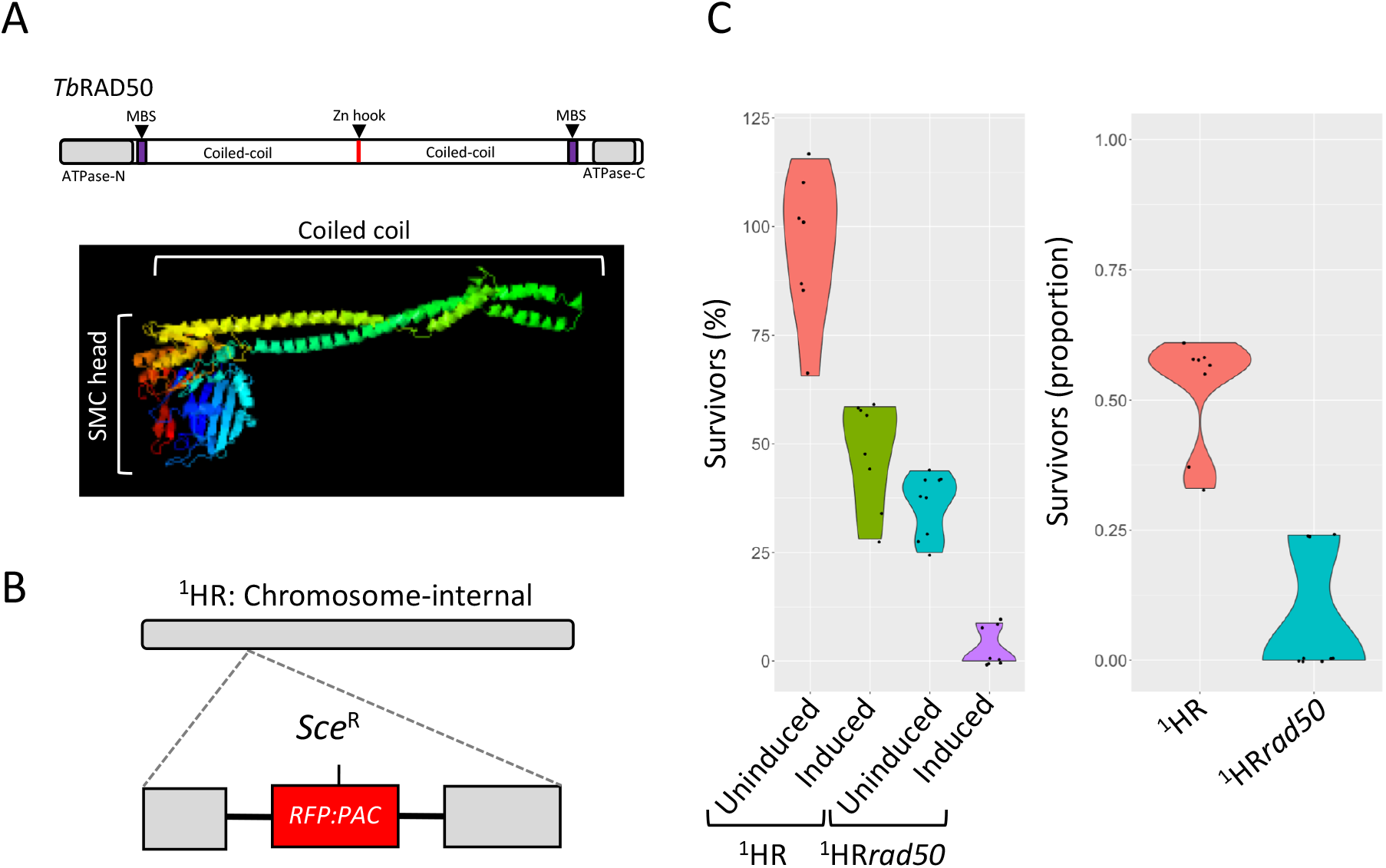
RAD50 is essential for DSB response and repair at a chromosome-internal locus. (A) Upper panel:Schematic of TbRAD50 with protein domains. Amino acid position of conserved domains are:ATPase - N, 4 – 170; MRE11 binding site (MBS), 182 – 205; Zn hook, 690 – 693; MRE11 binding site, 1158 – 1181; ATPase - C, 1243 – 1333. Lower panel:The structure of the *T. brucei* RAD50 was modelled using Phyre2 showing the SMC head domain with a coiled coil. (B) Schematic of the chromosome-internal DSB cell line with the I-SceI recognition site, Sce^R^, highlighted. (C) A clonogenic assay reveals survivors following a DSB at a chromosome-internal locus in the parental and ^1^HR*rad50* cell lines. Cells were plated out into media with or without tetracyline. The proportion of survivors was calculated by dividing the number of induced survivors by uninduced. R:P, red fluorescent protein:puromycin fusion gene. ^1^HR technical replicates; n=2, and with ^1^HR*rad50* biological replicates for the strains; n=2.

Clonogenic assays were plated out at either 32 cells per plate under both inducing and non-inducing conditions for ^1^HR and VSG^up^ strains and 480 cells per plate for VSG^up^ strains under inducing conditions. Plates were counted 5-6 days later and subclones selected for further analysis.

### Plasmid construction

For native *C*-terminal epitope tagging of Tb927.5.1700 / *RPA2* a 765-bp fragment was amplified using primers RPA28F:GATCAAGCTTATGGAAGGAAGTGGAAGTAA; and RPA28R:GATCTCTAGAAATGCCAAACTTACAATCATG and cloned in pNAT^xTAG 83^ using the HindIII and XbaI sites (underlined). The construct was linearized with XhoI prior to transfection. MRE11F5 (GATCgcggccgcATGGCCGAGAGGGCATC), MRE11R5 (GATCtctagaCAACGAAGATGTATGCCC), MRE11F3 (GATCgggcccCGATGGATAGTGGTAAT) and MRE11R3 (GATCggtaccCTAATAGTTATCTGGCA) were used to clone in target regions to generate pMRE11KOBLA and pMRE11KONEO. For transfection, 20 μg pMRE11KO *Blasticidin* (BLA) and *Neomycin* (NEO) plasmids were sequentially digested with Acc65I and NotI and cleaned by phenol-chloroform extraction and ethanol precipitation after each digestion. Strains were validated using MRE11F5 and MRE11R3 in a PCR assay. Heterozygous (+/−) and homozygous (−/−) knockout mutants of *RAD50* were generated by deleting most of replacing most of the gene’s open reading frame with either BLA and NEO. The strategy used is as described in ^84^; briefly, two modified versions of the plasmid pmtl23 were used to allow PCR-amplified 5’ and 3’ flanking untranslated regions of *RAD50* to be inserted around BLA and NEO cassettes (where the antibiotic resistance genes’ ORF were flanked by tubulin and actin intergenic regions). The selective drug markers, flanked by RAD50 5’ and 3’ untranslated regions, were then excised using NotI and transfected into *T. brucei*, and clones selected using 10 μg.ml^−1^ blasticidin or 5 μg.mL^−1^ G418.

### Immunofluorescence microscopy

Immunofluorescence analysis was carried out using standard protocols as described previously ^85^. Mouse α-Myc was used at 1:400 and rabbit α-γH2A ^55^ was used at 1:250. Fluorescein-conjugated goat α-rabbit and goat α-mouse secondary antibodies (Pierce) were used at 1:2000. Samples were mounted in Vectashield (Vector Laboratories) containing 4, 6-diamidino-2-phenylindole (DAPI). In *T. brucei*, DAPI-stained nuclear and mitochondrial DNA can be used as cytological markers for cell cycle stage ^86^; one nucleus and one kinetoplast (1N:1K) indicate G_1_, one nucleus and an elongated kinetoplast (1N:eK) indicate S phase, one nucleus and two kinetoplasts (1N:2K) indicate G_2_/M and two nuclei and two kinetoplasts (2N:2K) indicate post-mitosis. Images were captured using a ZEISS Imager 72 epifluorescence microscope with an Axiocam 506 mono camera and images were processed and in ImageJ.

### DNA analysis

Slot blots for detection of ssDNA were carried as described previously ^17^. ImageJ was used to generate linear density plots. The VSG probe was a 750 bp fragment *VSG2* fragment from a Pst1 digest of pNEG. The *RFP* probe was a 687-bp HindIII/NotI fragment encompassing the full ORF. Loading control was a 226 – bp product from *Tb427.01.570* (Dot1bKOF:TGGTCGGAAGTTGGATGTGA Dot1bKOR:CTTCCATGCATAACACGCGA).

PCR analysis of RAD50 nulls to confirms knock-out were done using standard PCR conditions with the following primers; a 402 bp product for RAD50 using RAD50KOF (CGTGAGAAACAGGAACAGCA) and RAD50KOR (AACACGTTTTTCCAACTCGG); a 399 bp product for *Blasticidin* ORF using BlaF (GATCGAATTCATGGCCAAGCCTTTGTCT) and BlaR (GATCCCATGGTTAGCCCTCCCACACATAA); and a 795 bp product for *Neomycin Phosphotransferase* ORF using NPTF (ATGATTGAACAAGATGGATTG) and NPTR (TCAGAAGAACTCGTCAAGAA). Analysis of subclones was previously described^21,27,35^ and used the following primers VSG221F (CTTCCAATCAGGAGGC), VSG221R (CGGCGACAACTGCAG), RFP (ATGGTGCGCTCCTCCAAGAAC), PAC (TCAGGCACCGGGCTTGC), ESAG1F (AATGGAAGAGCAAACTGATAGGTTGG), ESAG1R (GGCGGCCACTCCATTGTCTG), 2110X (GGGGTGAATGTTGGCTGTG), 2110Y (GGGATTCCCAGACCAATGA)

### VSG sequencing analysis

For the RT-PCR, the reaction mix were as following; 1 μg of cDNA, 1 x PCR buffer, 0.2 mM dNTPs, 1 μl each of SL (ACAGTTTCTGTACTATATTG) and SP6-14mer (GATTTAGGTGACACTATAGTGTTAAAATATATC) primers, H2O to 50 μl and 0.5 μl Phusion polymerase (New England Biolabs). For the PCR conditions. Five cycles were carried out at 94 °C for 30s, 50 °C for 30s and 72 °C for 2 min; followed by 18 cycles at 94 °C for 30s, 55 °C for 30s and 72 °C for 2 min. DNA concentration was measured using a Nanodrop. Libraries were prepared from *VSG* PCR products and sequenced on a BGI-Seq (BGI500) with a 150 bp paired-end read length with BGI Genomics Hong Kong.

Replicate libraries for WT uninduced, WT induced, VSG^UP^ uninduced and VSG^UP^ induced, VSG^UP^*rad50* uninduced and VSG^UP^*rad50* induced were sequenced on the BGIseq500 platform producing 8.03, 9.01, 7.60, 7.22, 6.66, 7.07, 6.90, 6.98 million reads per library, respectively. Reads were aligned to the *T. brucei* Lister 427 genome ^15^ with the cohort of minichromosomal VSGs added from the Lister 427 VSGnome ^13^ using bowtie2 ^87^ with the parameters --very-sensitive and BAM files created with samtools ^88^, aligning (WT uninduced) 97.76, 98.42, (WT induced) 97.69, (VSG^UP^*rad50* uninduced) 97.57, 98.60, (VSG^UP^*rad50* induced) 97.63, 97.80 percent of reads successfully. Reads counts per transcript were obtained using featureCounts^89^. Differential expression analysis was performed using EdgeR^90^on all genes, followed by filtering for *VSG* genes (1848 *VSG* sequences in total). An R script (https://github.com/LGloverTMB/DNA-repair-mutant-VSG-seq) was used to perform differential expression analysis, and generate Volcano and genome scale plots. BLAST analysis was performed locally using a database containing significantly up-regulated *VSG* genes from both conditions, including 2 kb of sequence upstream and downstream of the start and stop codons, respectively (except where sequences in the contigs 5’ or 3’ to the CDS were shorter than 2 kb excluding). For this analysis, minichromosomal *VSG* genes were excluded as the VSGnome does not contain any sequence beyond the CDS. The resulting database of 131 *VSGs* was queried using the *VSG2* sequence including 2 kb of sequence upstream of the CDS and all sequence between the stop codon and end of contig (1,272 nt). The BLASTn algorithm was used query the database using default parameters except allowing up-to 20 hits per subject sequence, and outputting up to 2,000 alignments. Alignments were filtered to remove overlapping hits from the same subject sequence using Microsoft Excel, retaining the one with the higher alignment score. Non-overlapping alignments were plotted using a custom R script (https://github.com/LGloverTMB/DNA-repair-mutant-VSG-seq). Lengths of average alignments were calculated for cohorts of VSGs up-regulated in both VSG^up^ and VSG^up^*rad50* or VSG^up^*rad50* only.

## RESULTS

### RAD50 is required for normal cell growth and DSB repair

RAD50, the largest component of the MRN complex, belongs to the structural maintenance of chromosomes (SMC) family of proteins ^48^ and has not been examined in *T. brucei*, though the gene has been reported to be essential in *Leishmania infantum ^49^*. The domain architecture of RAD50 is approximately palindromic (Figure 1A) and characterized by the presence of ATP-binding cassette (ABC)-ATPase domains at the N- and C-termini, each followed by an MRE11 binding site (MBS), and then by anti-parallel coiled-coil regions, which form linker structures that enable the MRN complex to act as a tethering scaffold to hold broken chromosomes together for repair ^50^. Between the antiparallel coiled-coils, a central Zn hook, a CxxC motif, facilitates Zn^2+^ dependent RAD50-RAD50 subunit interactions and is presumed to be important for tethering ^51^. A conformational change is invoked through binding of RAD50 to two ATP molecules, which then allows for binding to DNA^43^. Primary sequence comparison suggested all RAD50 domains are recognisably conserved in the putative *T. brucei* RAD50 homologue (Tb.927.11.8210; Supplementary Figure 1). Within the ATPase domains, the ABC nucleotide binding domain is defined by the conserved presence of Walker A, Q-loop, Signature, Walker B, D-loop, and H-loop motifs required to form the active ATPase site ^52^. Furthermore, structure prediction using Phyre^2 53^ modelled 503 residues (37 % of the sequence) of the *T. brucei* protein, revealing a SMC head domain and antiparallel coiled coil regions (Figure 1A).

To test the function of RAD50 in DSB repair, we used a previously validated *T. brucei* cell line, referred to as ^1^HR (Figure 1B), where a single I-*Sce*I meganuclease DSB can be induced in an *RFP-PAC* (red fluorescent protein – puromycin *N-*acetyltransferase) fusion cassette in the core region on chromosome 11 ^17^ (Figure 1B). We generated *rad50* null mutants (referred to as ^1^HR*rad50*) in these cells by sequentially replacing the two gene alleles with neomycin phosphotransferase (*NEO*) and blasticidin (*BLA*) resistance cassettes:PCR analysis of double antibiotic resistant clones confirmed *RAD50* loss and replacement (Supplementary Figure 2A and B) and demonstrates RAD50 is not essential in *T. brucei*. To determine the role RAD50 plays in DNA repair, we set up clonogenic assays. Cells were distributed across 96-wells plates under both I-*Sce*I non-inducing and inducing conditions, and wells with live cells scored after 5 - 7 days. This revealed a significant growth defect in the ^1^HR*rad50* null cells in unperturbed cells (Figure 1C Left panel and Table 1): 95 % of the WT ^1^HR cells survived compared with ~ 35 % of the ^1^HR*rad50* cells, revealing a 2.6 - fold decrease in cell survival. This growth impairment is likely due to the inability to repair spontaneous DSBs. Induction of the I-*Sce*I meganuclease results in ~ 95 % cutting and repair mainly by homologous recombination ^17^. Consistent with previous findings, in the WT ^1^HR strain ~ 48 % of cells are able to repair the DSB and survive (Figure 1C Left panel and ^17^ and Table 1), whereas in the ^1^HR*rad50* cells, a severe growth defect was seen following a DSB, with less than 3 % survival (a 16 - fold reduction), suggesting a significant defect in DSB repair (Figure 1C Left panel and Table 1). This was recapitulated when assessing the normalised survival efficiency (compared to uninduced survival) following an I-*Sce*I break (Figure 1C right panel and Table 1), indicating that a DSB is more lethal in the null mutant cells. In parallel we also tested the function of MRE11, as it forms a complex with RAD50, by generating null mutants through sequentially replacing the two gene alleles with *NEO* and *BLA* resistance cassettes; PCR analysis confirmed *MRE11* loss and replacement (Supplementary Figure 2C). The *mre11* nulls (referred to as ^1^HR*mre11*) also showed a growth defect cells in the unperturbed cells, with only 47 % of cells surviving cloning. Induction of the I-*Sce*I meganuclease resulted in a severe growth defect, with less than 2 % survival, suggesting a significant defect in DSB repair (Supplementary Figure 4A and Table 1) whose magnitude was very similar to ^1^HR*rad50* cells, which is expected given they have been shown to act in complex in other systems^37,38^.

**Table 1:**
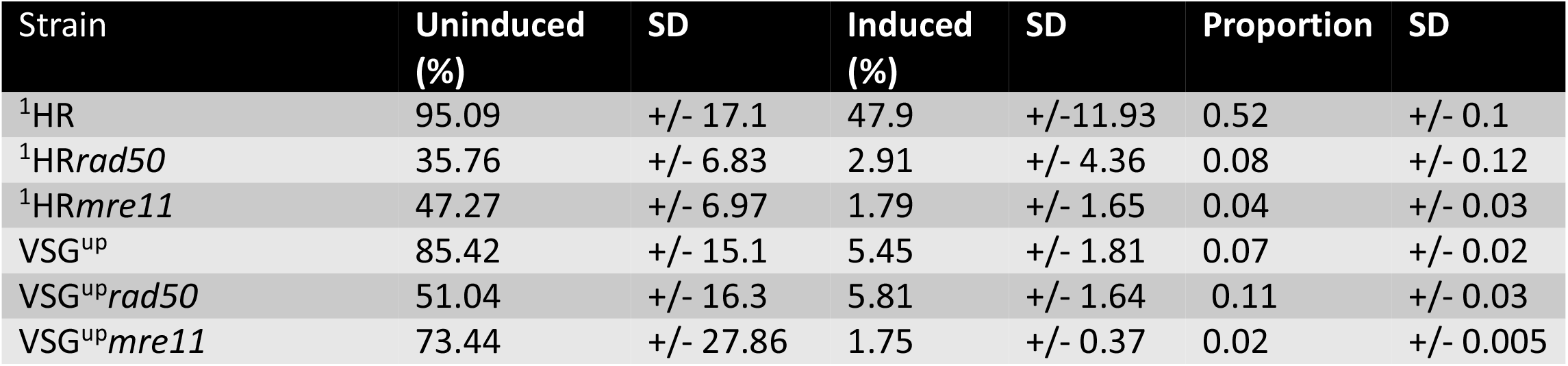
Details of clonogenic assays of strains described in this manuscript following a DSB.

We next asked what effect of the loss of RAD50 had on the mechanisms by which trypanosomes recognize a DSB lesion and initiate a signalling cascade resulting in DNA repair ^54^. In *T. brucei*, the DDR after an I-*SceI* induced DSB has been characterized thus far to include an increase of cells in G_2_/M ^17^, phosphorylation of histone H2A ^55^, break resection and accumulation of RAD51 foci at the site of the DSB ^17^. The cell cycle distribution of WT ^1^HR and ^1^HR*rad50* cells was assessed following induction of a DSB. In WT ^1^HR cells ~ 28% were in G_2_ 12 hours after I-*Sce*I induction and this returned to background levels (~15% of the population) by 24 hours. In contrast, no increase in G_2_ cells was seen after DSB induction in the ^1^HR*rad50* cells (Figure 2A), suggesting RAD50 is required for eliciting the G_2_/M checkpoint. In mammals the MRN complex recruits the ATM kinase to a DSB, where it phosphorylates H2AX ^37^.

**Figure 2:**
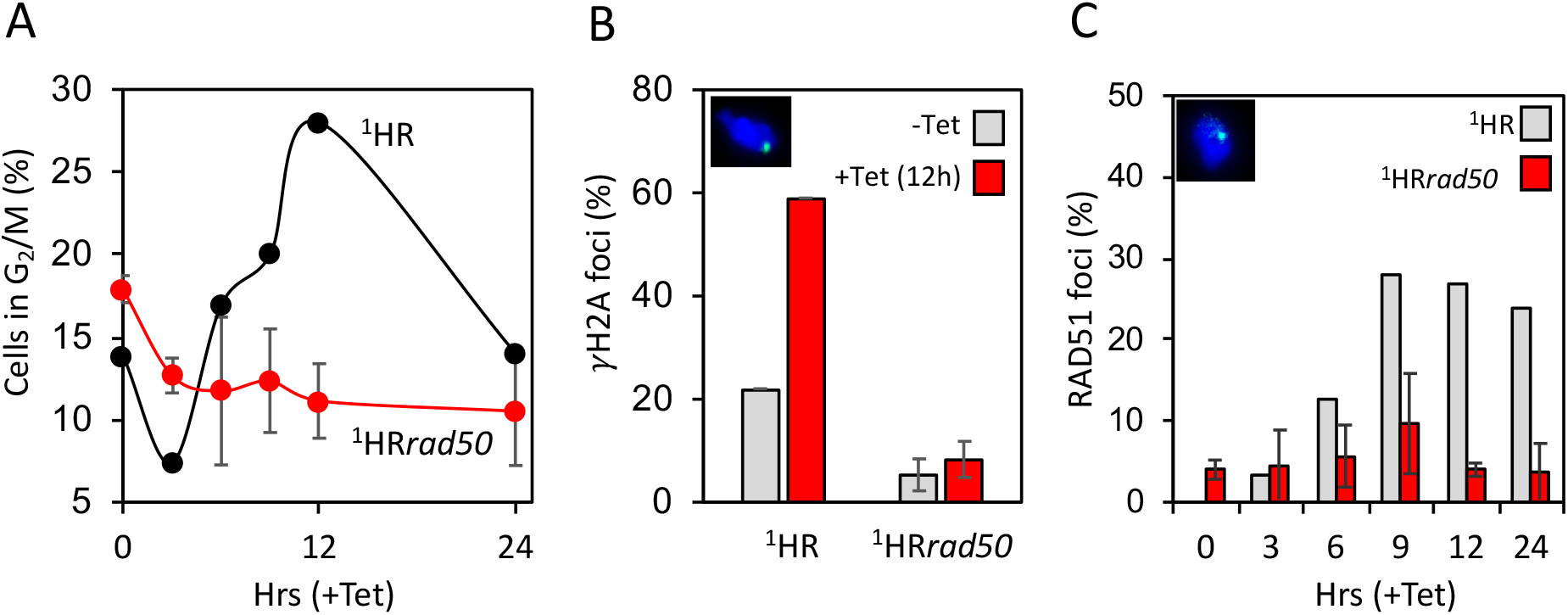
DNA damage response is compromised in ^1^HR*rad50* cells. (A) The number of cells in G2/M phase cells was counted by DAPI staining at several points following induction of an I-*Sce*I break in. G2 cells contain one nucleus and two kinetoplasts. (B) Immunofluorescence assay to monitoring γH2A foci. The number of positive nuclei were counted in uninduced cells and 12 hours post DSB. Inset showing a nucleus with a γH2A focus. n = 200 for each time point in the ^1^HR cell line and n= 400 for the ^1^HR*rad50* strain. Error bars, SD, for ^1^HR*rad50* biological replicates for the strains; n=2. (C) Immunofluorescence assay to monitoring RAD51 foci. The number of positive nuclei were counted in uninduced cells and 12 hours post DSB. Inset showing a nucleus, with a single RAD51 focus. n = 200 for each time point in the ^1^HR cell line and n= 400 for the ^1^HR*rad50* strain. Error bars, SD, for ^1^HR*rad50* biological replicates for the strains; n=2.

Using an antibody specific to the Thr130 phosphorylated form of *T. brucei* H2A, γH2A ^55^, we saw the expected background staining of ~15 – 20% of nuclei with foci in unperturbed WT ^1^HR cells, which increased to ~ 60% at 12 hours post I-*Sce*I induction (Figure 2B). In the ^1^HR*rad50* cells, the background level of γH2A foci was reduced to 5%, and the DSB-induced increase was drastically impaired (Figure 2B), with only 8% of cells containing γH2A foci. Repair at this locus is predominately via RAD51-dependent homologous recombination ^17^, and so we next assessed RAD51 foci assembly following DSB induction. In the WT ^1^HR strain, the number of detectable foci increased from 0 to 27% within 9 – 12 hours after I-*Sce*I induction. In contrast, in ^1^HR*rad50* cells the background level of RAD51 foci, before I-*Sce*I induction, was higher at 4%, and only increased to 10 % (~3 fold reduced) in response to a DSB (Figure 2C). Like the ^1^HR*rad50* cells, we detected fewer γH2A foci in ^1^HR*mre11* cells following a DSB (no foci detected, Supplementary Figure 4B) and a significant reduction in the number of RAD51 foci (14% compared with 36% in WT, Supplementary Figure 4B) and a loss of the G_2_/M checkpoint (8.5% compared with 28% in WT, Supplementary Figure 4B). These results reveal an important role for RAD50 and MRE11 in the DDR to a DSB in trypanosomes at a single copy locus and suggest wider roles in tackling spontaneous DNA damage.

### RAD50 restricts resection during allelic recombination

An early step in the DSB repair cycle is the formation of extensive 3’ ssDNA overhangs, initiated by MRE11 3’ – 5’ nuclease activity, which are a substrate for RAD51 nucleoprotein filament formation and act as a template for homology-directed repair ^37^. In light of the reduced accumulation of RAD51 foci after DSB induction in the absence of RAD50, we sought to determine whether the formation of ssDNA at the I-*Sce*I target locus was compromised, using slot blots. In the WT ^1^HR cells, ssDNA accumulated up to 12 h after I-*Sce*I induction and declined thereafter (Figure 3A), mirroring the phosphorylation of H2A and accumulation of RAD51 (Figure 2 B and C)^17^. Processing of the DSB in the ^1^H*Rrad50* cells appeared to be accelerated, with ssDNA signal peaking at 9 hours and declining thereafter (Figure 3A and Supplementary Figure 3). We conclude that DNA resection is not lost in the ^1^H*Rrad50* cells but the timing is affected, though we cannot say if the extent of resection is changed. Prior to RAD51 loading on to ssDNA, the trimeric RPA (replication protein A) complex binds the ssDNA and is subsequently displaced by RAD51 ^56^. Rescue of the *BLA* selectable marker in this strain (Supplementary Figure 2) allowed tagging of RPA2 with the myc epitope and subsequent localization. In WT ^1^HR cells the number of nuclei with RPA foci increased 5-fold (from 10% to 50%) following an I-*Sce*I break (Figure 3B). The ^1^HR*rad50* cells showed a pronounced increase in RPA foci prior to induction of a DSB, and only a marginal increase at 12 hours post DSB (~30% – 55%; Figure 3B). In the WT ^1^HR cells, a single RPA focus is most commonly seen in response to an I-*Sce*I break ^22^. However, we observed multiple RPA foci in both induced and uninduced ^1^HR*rad50* null cells (Figure 3C). We therefore tentatively conclude that most RPA signal in the ^1^HR*rad50* cells^22^ represents persistent, widespread damage, meaning it is unclear if loss of RAD50 alters the accumulation of RPA at the I-*Sce*I induced DSB in chromosome 1.

**Figure 3:**
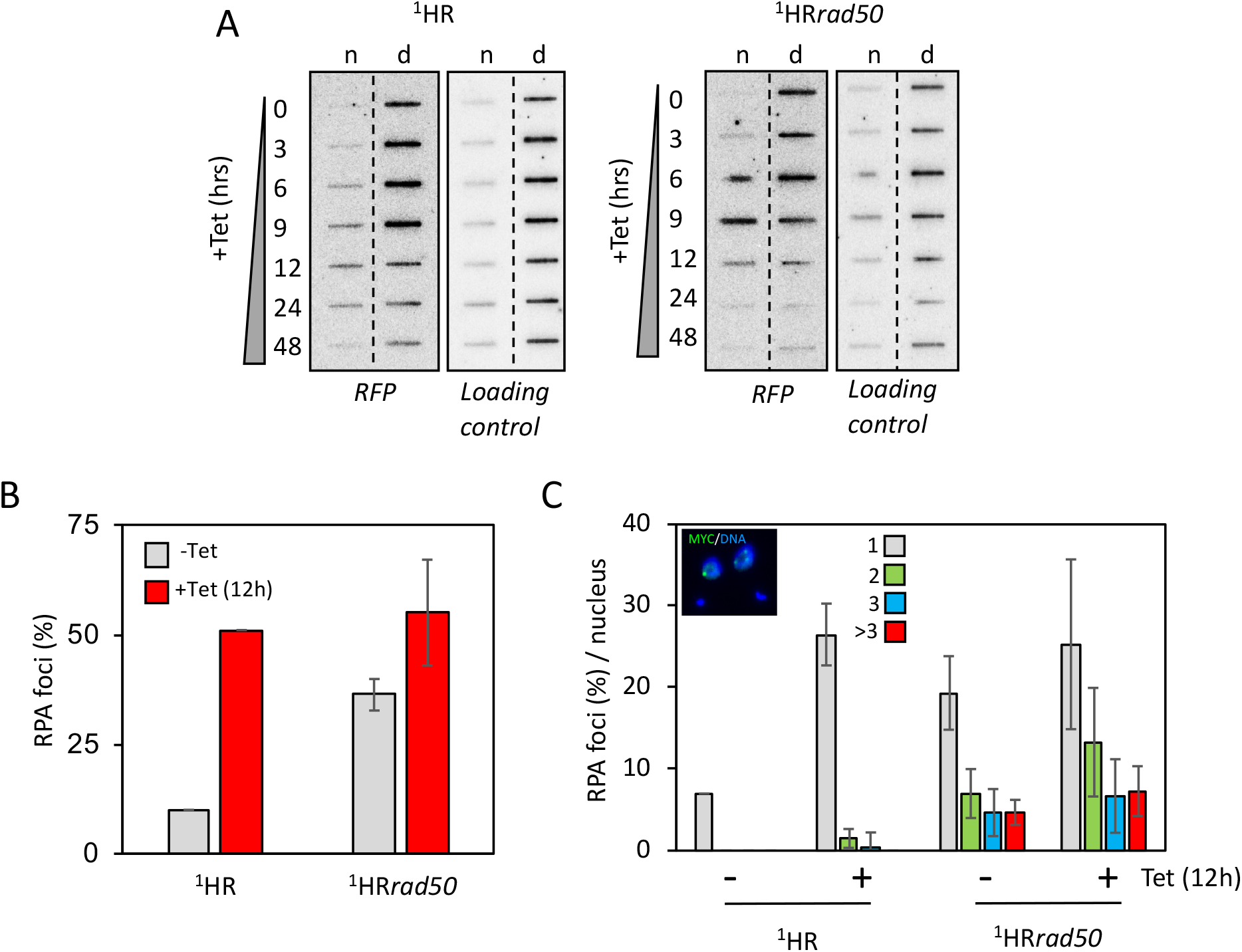
RAD50 directs resection at chromosome-internal locus. (A) Accumulation of ssDNA was monitored using slot-blots. Genomic DNA was extracted as indicated following I-*Sce*I induction. Ninety percent of the sample was ‘native’ (n; ssDNA) and ten percent was denatured (d). Probe *RFP* and the loading control are described in the materials and methods. (B) Immunofluorescence assay to monitoring RPA foci. The numbers of positive nuclei were counted in uninduced cells and 12 hours post DSB. n = 200 for each time point in the ^1^HR cell line and n=400 for the ^1^HR*rad50* strain. Error bars, SD, for ^1^HR*rad50* biological replicates for the strains; n=2. (C) Immunofluorescence assay to monitor the number of RPA foci per nucleus. The number of RPA foci was counted in uninduced cells and 12 hours post DSB. n = 200 nuclei for each time point in the ^1^HR cell line and n= 400 nuclei for the ^1^HR*rad50* cells. Inset showing representative nuclei, with RPA foci. Error bars, SD, for ^1^HR*rad50* biological replicates for the strains; n=2.

### RAD50 is crucial for homologous recombination in *T. brucei*

To explore how trypanosomes repair a DSB in the ^1^HR*rad50* nulls, DSB-survivors from the clonogenic assay were scored for repair by homologous recombination or MMEJ by a PCR assay (Figure 4A) using sets of primers that flanked the *RFP-PAC* cassette. In the surviving subclones, sensitivity to puromycin is indicative of cleavage by I-*Sce*I ^17^. All twelve of the surviving subclones were sensitive to puromycin, indicating cleavage by I-*Sce*I ^17^ and disruption of the *RFP:PAC* cassette (data not shown). Eleven of these clones showed repair by MMEJ, as seen by the reduction in the size of the PCR product as compared to the controls (Figure 4B, RFP and PAC primer pair^17^), or loss of the entire cassette (Figure 4C, 2110X and 2110Y primer pair ^57^). Sequencing revealed repair by MMEJ using the *Xcm*1 sites that flanked the *RFP-PAC* cassette in clones 12, 15, 16 and 17 (Figure 4C lower panel). Sequencing of the 2110X-Y product in clone 8 revealed repair using the homologous template, suggesting homologous recombination can still occur, although is significantly impaired. In the *mre11* nulls, 14 out of 15 clones repaired by MMEJ (Supplementary Figure 4C, clone 10, 17 and 19 show a PCR product of reduced size and for the remaining clones the 2110XY PCR product was sequenced revealing 11 clones had repaired by MMEJ). These data show a significant shift in the pathway used to repair a DSB in the ^1s^HR*rad50* and ^1^HR*mre11* null cells at a chromosome-internal locus, with repair by MMEJ dominating (Figure 4D), compared with the pronounced predominance of homologous recombination in WT ^1^HR cells ^17^.

**Figure 4:**
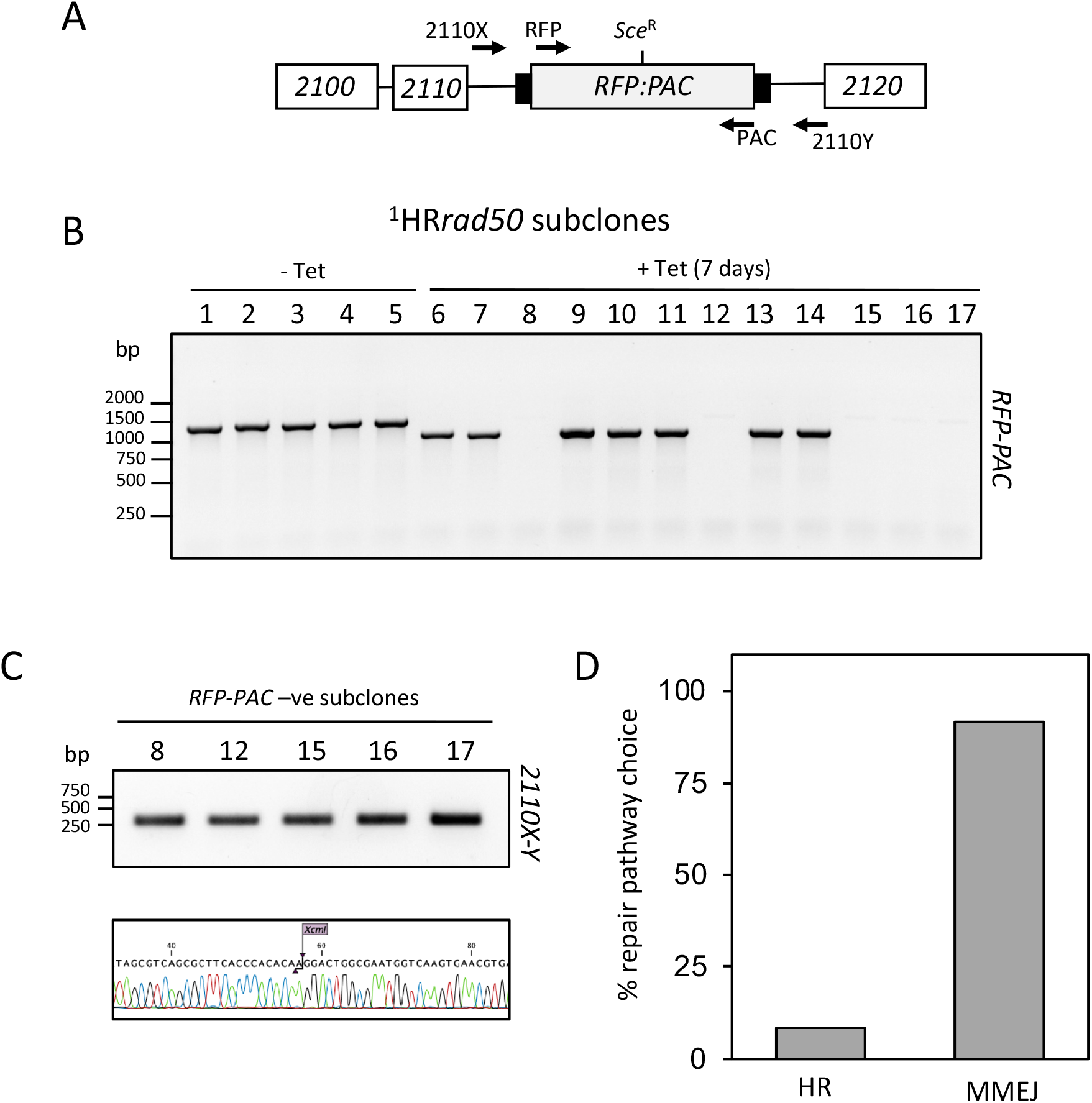
RAD50 is required for homologous recombination. PCR analysis of ^1^HR repaired subclones. (A) Schematic showing the 2110 locus and position of the *Sce* recognition site (Sce^R^). Position of primers indicated by arrows. Primer sequence detailed in materials and methods. (B). PCR assay of repaired subclones showing *RFP:PAC* presence or absence. (C) Upper panel: PCR assay of repaired subclones that were negative for *RFP:PAC*. Lower panel: Sanger Sequence trace showing XcmI site. (D) Percentage of survivors for each repair pathway choice. n= 12 clones. Arrows indicate position of primers. White box, genes; Grey box, *RFP – PAC* fusion gene; black box, UTRs.

### Loss of RAD50 increases survival following a DSB at the active VSG-ES

Trypanosomes rely on homologous recombination to facilitate antigenic variation. We therefore wanted to test the role of RAD50 in *VSG* switching. We generated *RAD50* nulls in a cell line where the I-*Sce*I recognition site is fused to a *puromycin* selectable marker and inserted immediately downstream of the major block of 70-bp repeats and upstream of *VSG2* in Bloodstream form Expression site 1 (BES1) (on chromosome 6a), the active *VSG*-ES in this strain (Figure 5A). The resulting cell line is known as VSG^up 27^. Cell survival following a DSB at this position is contingent upon VSG switching, most commonly using the 70-bp repeats and replacing the active *VSG* via break-induced replication ^26,27^. We generated *rad50* null VSG^up^ strains (VSG^up^*rad50*), by replacing the two gene alleles with *NEO* and *BLA* resistance cassettes: PCR analysis of double antibiotic resistant clones confirmed *RAD50* loss and replacement (Supplementary Figure 2A and B). Using a clonogenic assay we found that, like in the ^1^HR strain, there was a growth defect (1.6-fold reduction) in the VSG^up^*rad50* nulls compared with VSG^up^ WT cells in the absence of I-*Sce*I induced damage (Figure 5B; left panel and Table 1). This effect again suggests an impaired ability to repair spontaneous damage within the mutant cell. However, quite differently to ^1^HR cells, following induction of an I-*Sce*I DSB we observed an increase in survival in the VSG^up^*rad50* nulls compared with induced VSG^up^ WT cells (Figure 5B; left panel and Table 1). These data indicate that RAD50 suppresses DSB repair at a *VSG*-ES, the opposite of its role at a chromosome-internal DSB. In contrast, survival is reduced in the VSG^up^ *mre11* nulls relative to VSG^up^ WT, with only 1.75% able to survive a DSB (Supplementary Figure 6A and Table 1). We then assessed the DDR in the VSG^up^ cell line. As in the ^1^HR cells, following a DSB, the number of cells that accumulate in G_2_/M increases to ~ 30 % in the VSG^up^ WT cell line ^27^, and this cell cycle checkpoint was lost in the VSG^up^*rad50* cells (Figure 5C) and in the VSG^up^*mre11* cells (Supplementary Figure 6B). In the VSG^up^ WT cell line, the number of γH2A foci increased from 20% to 41% after I-*Sce*I induction, as has been previously reported ^27^. In the VSG^up^*rad50* nulls, γH2A foci were only detected in 3% of uninduced cells, and this increased to 11% following a DSB (Figure 5D). A similar phenotype was seen in the VSG^up^*mre11* cells, with the number of γH2A foci increasing from 1.3% to 7% (Supplementary Figure 6B). Thus, while it appears that loss of RAD50 or MRE11 diminishes the capacity of cells to phosphorylate H2A in response to a DSB at this locus, the increased survival in the VSG^up^*rad50* cells suggests that while it is required for an efficient DDR, in its absence the cells are more adept at repair.

**Figure 5:**
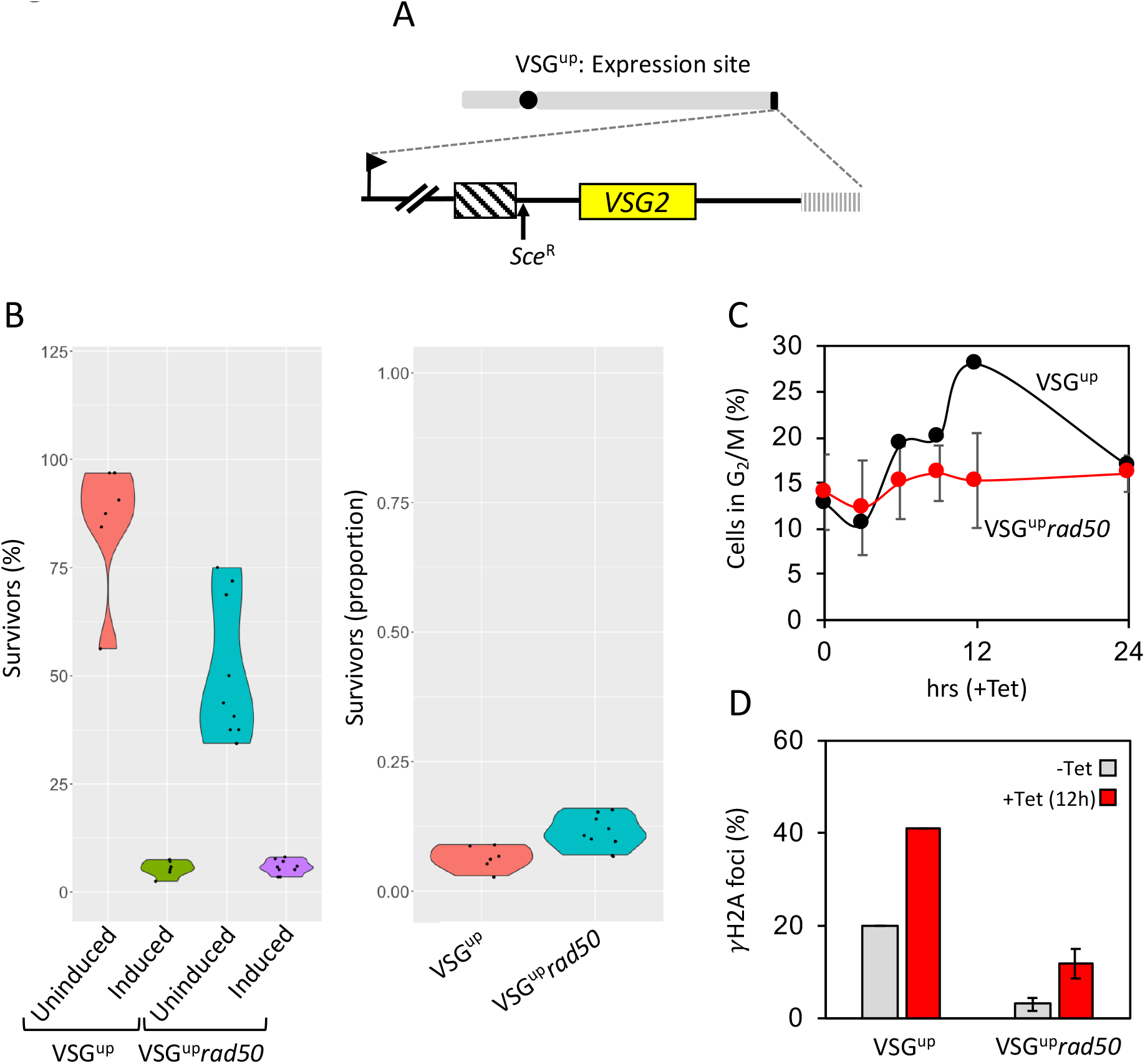
RAD50 suppresses repair at a subtelomeric locus. (A) A schematic of the active expression site DSB cell line with the I-*Sce*I recognition site, *Sce*^R^, highlighted. (B) A clonogenic assay reveals the survivors following a DSB at the active expression site in the parental and VSG^UP^*rad50* cell lines. Cells were plated out into media with or without tetracyline and counted after seven days. Other details as in Figure 1. (C) The number of cells in G2/M phase cells was counted by DAPI staining at 0 hours and 12 hours following induction of an I-*Sce*I break in. G2 cells contain one nucleus and two kinetoplasts. (D) Immunofluorescence assay to monitoring γH2A foci. The nuclei with γH2A foci were counted in uninduced cells and 12 hours post DSB. n = 200 for each time point in the VSG^up^ cell line and n= 400 for the *RAD50* null. Error bars, SD, for VSG^up^*rad50* biological replicates for the strains; n=2. Arrow, RNA Pol 1 promoter; box with diagonal lines, 70-bp repeats; vertical lines, telomere.

Using a series of assays (Figure 6A and Supplementary figure 5) we next looked at DNA rearrangements in the *VSG*-ES to determine how the VSG^up^*rad50* cells repair an induced DSB. 25 subclones were selected from the clonogenic assay and tested for sensitivity to puromycin, asking about the frequency of loss of the puromycin gene from the *VSG*-ES. 24 out of 25 subclones were sensitive to 1 μg.ml^−1^ puromycin, indicating that the majority of the population had been subject to a DSB and had deleted the resistance cassette. Immunofluorescence using antibodies against VSG2, the VSG expressed from the modified *VSG*-ES, showed that 23 out of 24 puromycin sensitive subclones had switched VSG (Figure 6B). This is comparable to what is seen in the VSG^up^ WT strain, suggesting loss of RAD50 does not affect the cell’s ability to undergo VSG switching. The single puromycin resistant clone was VSG2 positive, suggesting I-*See*I did not cut (Supplementary Figure 5, subclone 14).

**Figure 6:**
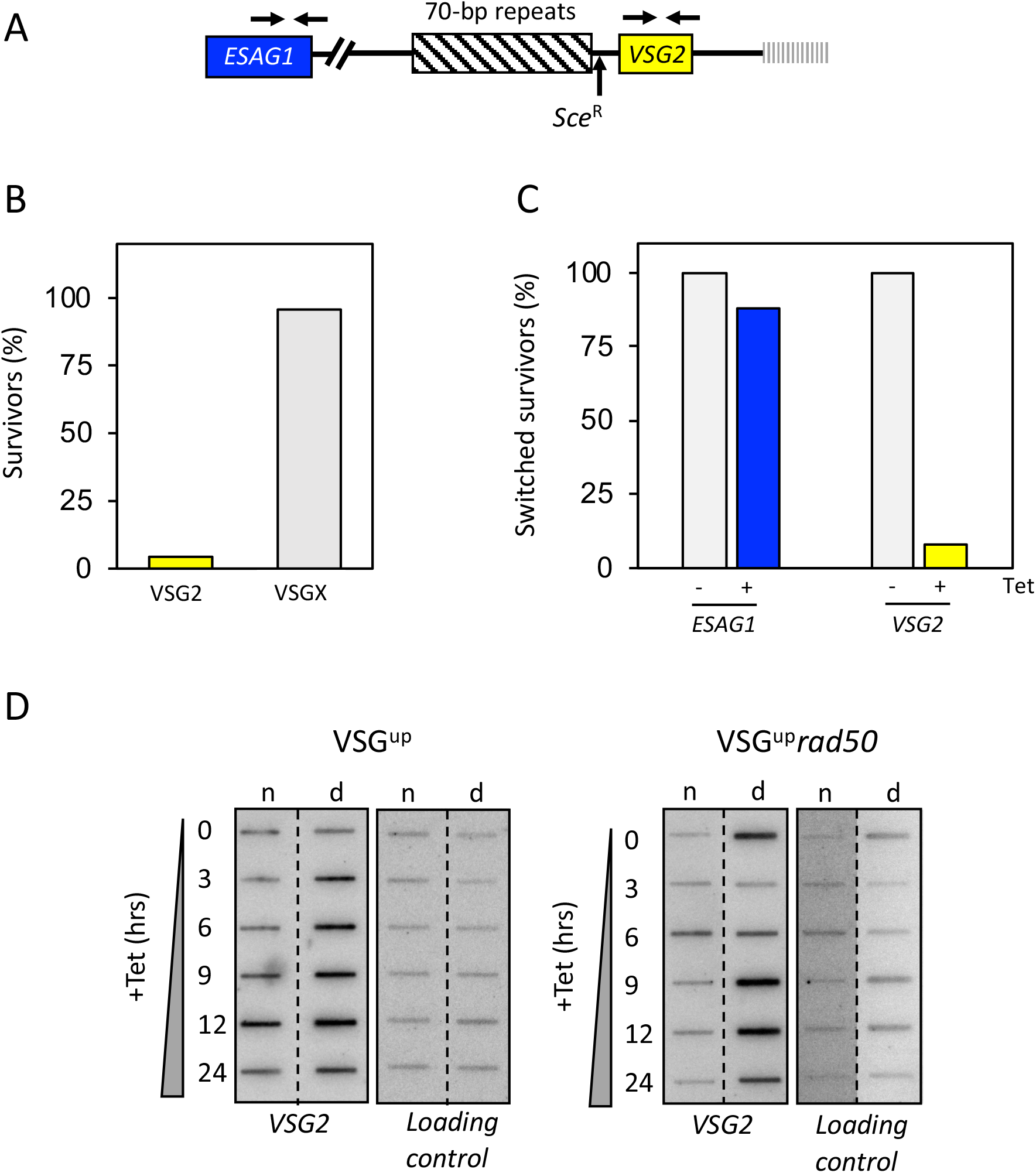
RAD50 is not required for VSG switching. (A) A schematic map shows the primer position at the active expression site. (B) Immunofluorescence assay for VSG2, showing the percentage of switched survivors in the VSG^up^*rad50* cell line. (C) PCR analysis shows the percentage of switched survivors that kept *ESAG1* and *VSG2*. n= 24 clones. Arrows indicate position of primers; box with diagonal lines, 70-bp repeats; vertical lines, telomere. (D) Accumulation of ssDNA was monitored using slot-blots. Genomic DNA was extracted as indicated following I-*Sce*I induction. Ninety percent the sample was ‘native’ (n; ssDNA) and ten percent was denatured (d). Probe *VSG2* and the loading control are described in the materials and methods.

We then looked at DNA rearrangements in the ES using primers specific to *VSG2* and *ESAG1*. *ESAG1* is found upstream of a block of 70-bp repeats in the active *VSG*-ES and cells that retain *ESAG1* are presumed to have repaired by gene conversion using the 70-bp repeats. Both *VSG2* and *ESAG1* were retained in five out of five uninduced control subclones (Figure 6C and Supplementary Figure 3). Of the 24 puromycin sensitive subclones (i.e. cleaved by l-*See*l), *ESAG1* was retained in 23 and *VSG2* was lost in 23 (Figure 6C and Supplementary Figure 5).

One subclone was found to be VSG2 negative by immunofluorescence, puromycin sensitive, and *ESAG1* positive and *VSG2* positive by PCR. This suggests that this single clone had switched by *in situ* transcriptional activation of another *VSG*-ES (Figure 6B and C and Supplementary Figure 3), whereas all other clones had undergone VSG switching by recombination. In the VSG^up^*mre11* cells, all the puromycin sensitive subclones had switched VSG and lost the *VSG2* gene as seen by PCR analysis (Supplementary Figure 4C). 18 out of the 20 subclones had also retained *ESAG1*. As with the VSG^up^*rad50* cells, these clones are presumed to have repaired by gene conversion using the 70-bp repeats. We then assessed the formation of ssDNA and found that as in the ^1^HR*rad50* nulls, the formation of ssDNA was not abolished but seemed to accumulate earlier - here at 6 hours post DSB induction (Figure 6D). These data suggest that loss of RAD50 or MRE11 does not impair the cell’s ability to undergo switching by DNA recombination.

### RAD50 promotes recombination using long stretches of homology

Increased survival of the VSG^up^*rad50* nulls after induction of a DSB suggested a hyper-recombinogenic mechanism in this locus, through which the cells are able to repair and switch at a higher rate than in the parental cell line. One explanation for such enhanced repair could be through greater access to the silent *VSG* repertoire. In the *T. brucei* genome there are in excess of 2000 VSG genes found at the subtelomeric arrays^16,13,15,58^, 90% of which are associated with a stretch of 70-bp repeat sequence^14^ that can be used for homology during repair^9^. To ask if differences in VSG repertoire access explains increased survival of VSG^up^*rad50* cells after DSB induction, we used VSG-seq ^10^. In the VSG^up^ WT cell line, 83 *VSG* gene transcripts were significantly enriched in the induced cells compared with uninduced (Figure 7A) (greater than 2 log_2_ fold change and *p* value of less than 0.05). In the VSG^up^*rad50* cell line, a greater number of *VSGs* were detected: here 225 *VSG* transcripts were significantly enriched after I-*Sce*I induction (Figure 7A). To understand this increase, we then looked at the genomic position of the *VSG* cohorts. In the VSG^up^ cells, we found that approximately equal numbers of enriched *VSGs* mapped to the *VSG*-ES and minichromosomes relative the megabase arrays, despite the much greater number of genes in the latter component of the archive (Figure 7B, Supplementary Figure 7A). These data appear consistent with VSG switching having followed a loose hierarchy, as previously published ^59^, with telomeric *VSG* preferred as donors. In the VSG^up^*rad50* cell line, enriched *VSG* genes also mapped to the *VSG*-ESs, minichromosomes and megabase arrays, but a significantly higher proportion (67% compared with 34% in WT) were from subtelomeric arrays (Figure 7B, Supplementary Figure 7A). This suggests that the VSG^up^*rad50* cells are able to access a greater proportion of the silent *VSG* archive for repair and VSG switching. To ask if increased VSG switching in the absence of RAD50 could be explained by changes in the mechanism of recombination, we looked at the length of homology used for repair. Using BLAST analysis, we queried the significantly enriched *VSGs* against the telomeric end of BES1, searching for regions of homology (Figure 7C and Supplementary Figure 7B). This analysis revealed that, compared with the VSG^up^ WT cells, *VSG* genes activated in the VSG^up^*rad50* cells shared shorter stretches of homology with the active *VSG2* locus.

**Figure 7:**
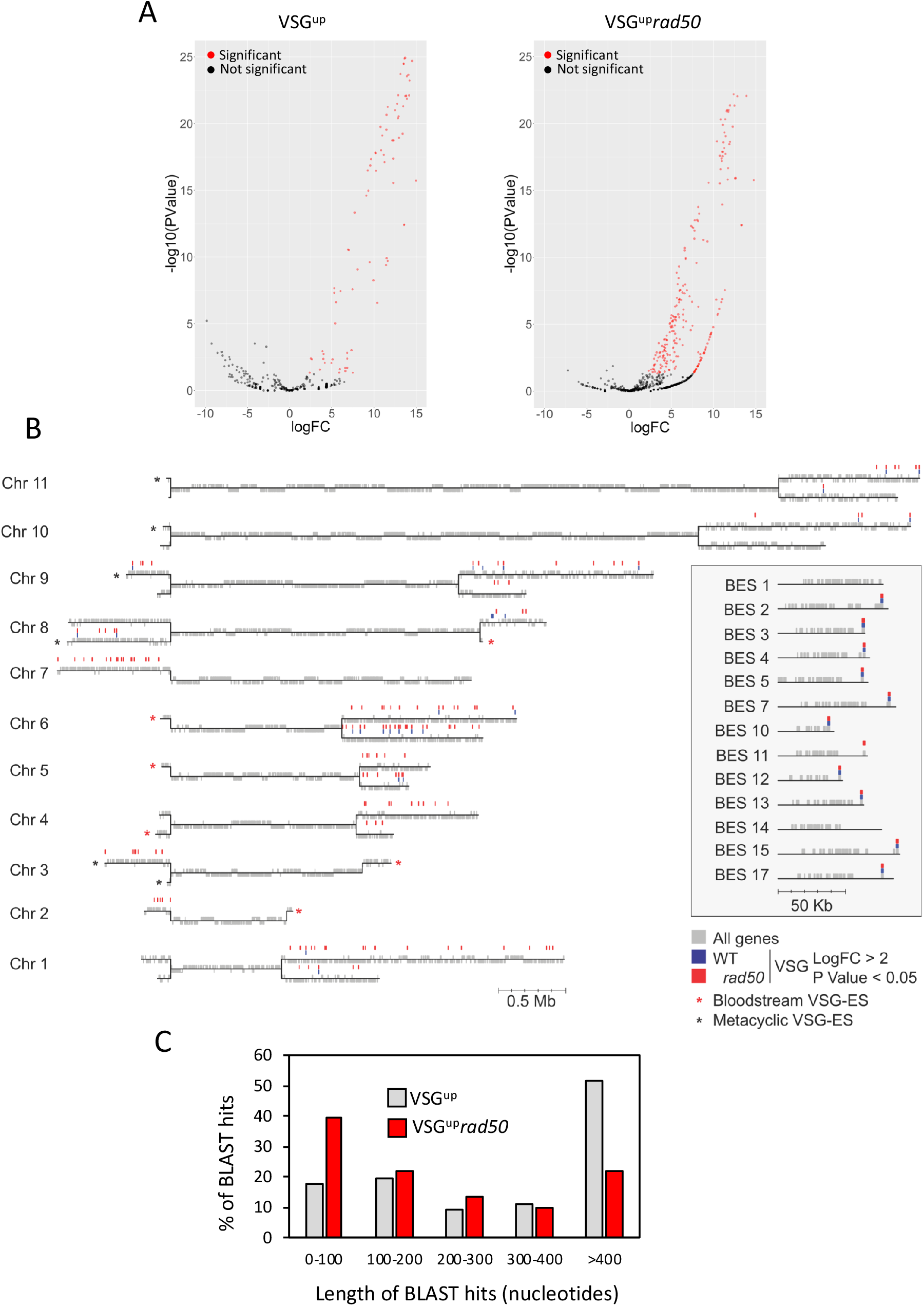
RAD50 restricts antigenic variation. (A) Volcano plots of VSG-seq showing log_2_ fold change vs. log_10_ P value for VSG^up^ and VSG^up^*rad50* strains. Red genes are significantly up-regulated (P value < 0.05 and log_2_ FC > 2. (B) *T. brucei* 427 Genome map showing all 11 megabase chromosomes in black lines and all genes in grey. Significantly up-regulated *VSG* genes from either VSG^up^ (blue) or VSG^up^*rad50* (red). Inset box shows all 13 BES with significantly up-regulated *VSG* genes from either VSG^up^ (blue) or VSG^up^*rad50* (red). (C) Chromosome 6 map showing all genes in grey and significantly up-regulated *VSG* genes from either VSG^up^ (blue) or VSG^up^*rad50* (red). (C) Percentage usage for the length of the BLAST hits from either VSG^up^ (blue) or VSG^up^*RAD50* (red).Box with diagonal lines, 70-bp repeats; vertical lines, telomere; grey box, 3’UTR

The most commonly used length of homology in the VSG^up^*rad50* cells was 100 bp (40% of survivors), whereas the VSG^up^ WT cells most frequently used >400 bp (51% of survivors) (Figure 7C). Thus, increased survival in the absence of RAD50 is due to increased access to archival VSGs due to more frequent DSB repair using short stretches of homology.

## DISCUSSION

Central to the DDR and subsequent repair is the MRN complex, within which the MRE11 – RAD50 heterodimer forms the catalytic core. In fact, this core complex is so fundamental to repair that it is conserved across bacteriophages, bacteria, archaea and eukaryotes ^60^. Kinetoplastids rely on homology-based recombination as the major form of repair, since no data has suggested the use of non-homologous end-joining after I-*Sce*I induction of a DSB at both at chromosome-internal^17^ and telomeric loci in *T. brucei*, or after CRISPR-Cas9 DSB formation in *Leishmania*, *T. cruzi* or *T. brucei ^61–63^*. At least in *T. brucei*, HR predominates over MMEJ after induction of a DSB^17^, and all evidence suggests RAD51-dependent HR dominates as the major form of repair reaction during VSG switching at the *VSG*-ES ^19^. What dictates the preference of HR over MMEJ in these reactions in *T. brucei* and related kinetoplastids is unknown. Here, we have shown that *T. brucei* RAD50 is critical for normal cell growth and efficient homology-based DNA repair after induction of a DSB, which is consistent with a conserved role in DSB recognition and repair. We demonstrate that in the absence of RAD50 the DDR is severely compromised, as evidenced by the loss of γH2A accumulation and the G_2_/M cell-cycle checkpoint at both subtelomeric and chromosome-internal DSBs, and the RPA focal accumulation. We also show that HR is dependent on RAD50, as null mutants displayed compromised RAD51 foci formation and switched repair pathway choice from HR to MMEJ at a chromosome-internal DSB. In contrast, at a *VSG*-ES we demonstrate that RAD50 restrains the DDR, since in its absence break processing dynamics were altered, cell survival improved and a greater diversity of VSGs were activated following a DSB. Thus, this work provides insight into DSB repair pathway choice, compartmentalization of repair with specific relevance for understanding trypanosome antigenic variation and genome maintenance in all kinetoplastids.

In *T. brucei* RAD51-dependent HR accounts for approximately 95% of DSB repair at a chromosome-internal locus, and a DSB triggers a conventional DDR^17^. The data shown here indicate that the role of *T. brucei* RAD50 in recognizing a DSB and initiating the DDR appears consistent with its role in other eukaryotes^64,65^. Previous characterization of MRE11 has revealed a role in HR and genomic stability in both *T. brucei* ^46,47^ and *Leishmania* ^49,66^, suggesting RAD50 and MRE11 act together. Here we show that that MRE11 is essential for the DDR and HR and, in its absence, repair is predominantly achieved by MMEJ. We expect that the reduced survival of ^1^HR*rad50* and VSG^up^*rad50* cells is due to unrepaired DNA damage that results in cell death, which is also seen in *mre11* mutants ^46,47^. The persistence of RPA foci in the ^1^HR*rad50* cells is further evidence of defective repair in these cells, explained by RPA remaining associated with ssDNA at sites of damage. This observation may also reflect a damage tolerance strategy employed by *T. brucei* ^22^, but in these circumstances γH2A and RAD51 are not recruited to the break after replication, unlike what is seen in *T. cruzi* where ionizing radiation tolerance is dependent on RAD51^67^. DNA resection following a DSB is one of the early steps in the DDR. MRN is an early acting complex in DSB processing, initiating resection^39,68^ and recruiting the SGS2 nuclease and DNA2 and EXO1 for long-range resection thereafter^39,69^. The lack of resection defect in the ^1^HR*rad50* mutants, coupled with the altered dynamics of the formation and resolution of ssDNA, suggests that in trypanosomes there is flexibility in the nucleases able to initiate and resect across a DSB and, in the absence of MRN, unknown nucleases can act in an unrestricted manner that exceeds the resection rate of WT cells.

Loss of RAD50 in *T. brucei* reveals a striking difference in the response to a DSB at a chromosome-internal location and in the active *VSG*-ES, with dramatically impaired survival at the former and improved survival at the latter. This locus-specific effect of *rad50* nulls was not reproduced in the *mre11* nulls. It is possible that this difference arises from a shared change in DSB repair reaction mechanism due to loss of the MRN complex. After DSB induction, loss of RAD50 results in lower levels of yH2A and RAD51 foci, allied to more rapid formation and resolution of ssDNA at a chromosome-internal site, with the result that DSB repair no longer favours HR but instead mainly uses MMEJ. An explanation for these effects could be that loss of RAD50 leads to impaired DSB processing by MRE11, allowing altered resection that leads to lowered levels of RAD51 nucleoprotein filament formation and increased recruitment of the unknown *T. brucei* machinery for MMEJ. If the MMEJ reaction was more efficient than HR, the more rapid resolution of ssDNA would be explained. If this same redirection of DSB repair occurred in the *VSG*-ES, why might the difference in survival occur? A simple explanation might be that MMEJ repair is more error-prone (such as by mutation or translocation) than HR^70^, and the use of error-prone MMEJ has a greater impact in the chromosome core, where it would affect housekeeping genes, than in the subtelomeres, where there is a huge abundance of silent VSG substrates. Because the machinery that directs MMEJ in *T. brucei* or any kinetoplastid has not been characterised, directs tests of this suggestion are not yet possible. In addition, it remains to be shown that MMEJ directs the activation of silent array *VSGs* after a DSB in the *VSG*-ES, since the length of homologies we describe here (predominantly ~100 bp) appear longer than the ~5-15 bp microhomologies so far detailed in experimental characterization of *T. brucei* MMEJ^21,71^. Nonetheless, previous work has shown considerable flexibility in the substrate requirements of recombination in *T. brucei^71^*, and linked MMEJ and HR-mediated repair of a DSB has been detailed^21^. Thus, it remains possible that MRN is pivotal in determining the routing of DNA repair throughout the *T. brucei* genome.

DNA DSBs are a potent trigger for *T. brucei* antigenic variation and it has been shown that the subtelomeric *VSG*-ES are fragile and prone to DSBs ^27^. A DSB upstream of the active *VSG* triggers antigenic variation, and subsequent repair by recombination is facilitated by stretches of 70-bp repeat that provide homology^9,26,27^, via either a RAD51-dependent or independent pathway^27^. Given this, it remains perplexing that mutation of MRE11 was not previously shown to alter the rate of VSG switching^46^, unlike the clear alteration in survival and repair pathway seen here after loss of RAD50 and induction of a *VSG*-ES DSB. These data may suggest an unconventional signalling pathway following a DSB at the active *VSG*-ES that allows the cells to bypass the conventional MRN / ATM pathway. One way that might occur is that the DSBs in the *VSG*-ES do not arise directly, but are generated by routes such as clashes between replication and transcription ^31^, or the formation of RNA-DNA hybrids^72–74^, which might not elicit recruitment of MRN or ATM^75,76^. Our data reveals that increased survival in the VSG^up^*rad50* cells is due to their ability to access more of the genomic *VSG* archive for repair, and the repair reaction occurs as a result of recombination using shorter stretches of homology. RAD50, therefore, appears to suppress antigenic variation by committing *T. brucei* to RAD51-dependent repair pathway that is reliant on long stretches of donor homology. We also note that we observe a striking number of donor *VSG* genes used for repair on Chromosome 6 – BES1, which contains the I-SceI site is found on Chromosome 6a. Indeed, several other factors have also been shown to suppress antigenic variation: HAT3^35^, TOPO3α^32^ and the telomere-interacting factor TIF2^77^. Antigenic variation following a DSB is driven by break-induced repair^26,27^ and, in yeast, RAD51-independent BIR has been shown require as little as 30 bp of homology for repair ^78^. Thus, loss of RAD50 may allow *T. brucei* to enlist RAD51-independent BIR to respond to a DSB in the *VSG*-ES, using shorter stretches of homology, rather than MMEJ. Irrespective, the question arises as to why *T. brucei* might limit the range of VSGs that can be used in antigenic variation, since this appears counter to the observed diversity of VSGs seen during infections^10,11^. It is possible that RAD50-mediated restraint of *VSG* recombination is needed to preserve the *VSG* archive, saving genes for use in prolonged infections. A more radical possibility is that RAD50/MRN control underlies the hierarchy of VSG expression, directing DSB repair to telomeric *VSG* substrates early an infection and then giving way to a distinctly signalled reaction, such as MMEJ, that allows access to the whole *VSG* archive later.

## DATA AVAILABILITY

All script is hosted on GitHub: https://github.com/LGloverTMB/DNA-repair-mutant-VSG-seq All data has been deposited onto the ENA under study accession number: PRJEB37290 and unique study name: ena-STUDY-INSTITUT PASTEUR-15-03-2020-20:11:26:661-115

## ACKNOWLEDGEMENTS

A-K.M, M.P, A.D-H and L.G performed the experiments, S.H performed the VSG-Seq data analysis, R.M and L.G discussed the results and wrote the manuscript. A-K.M, S.H, R. M and L.G edited the manuscript. We would also like to that David Horn for discussions and support.

## TABLE AND SUPPLEMENTARY FIGURE LEGENDS

**Supplementary figure 1:**
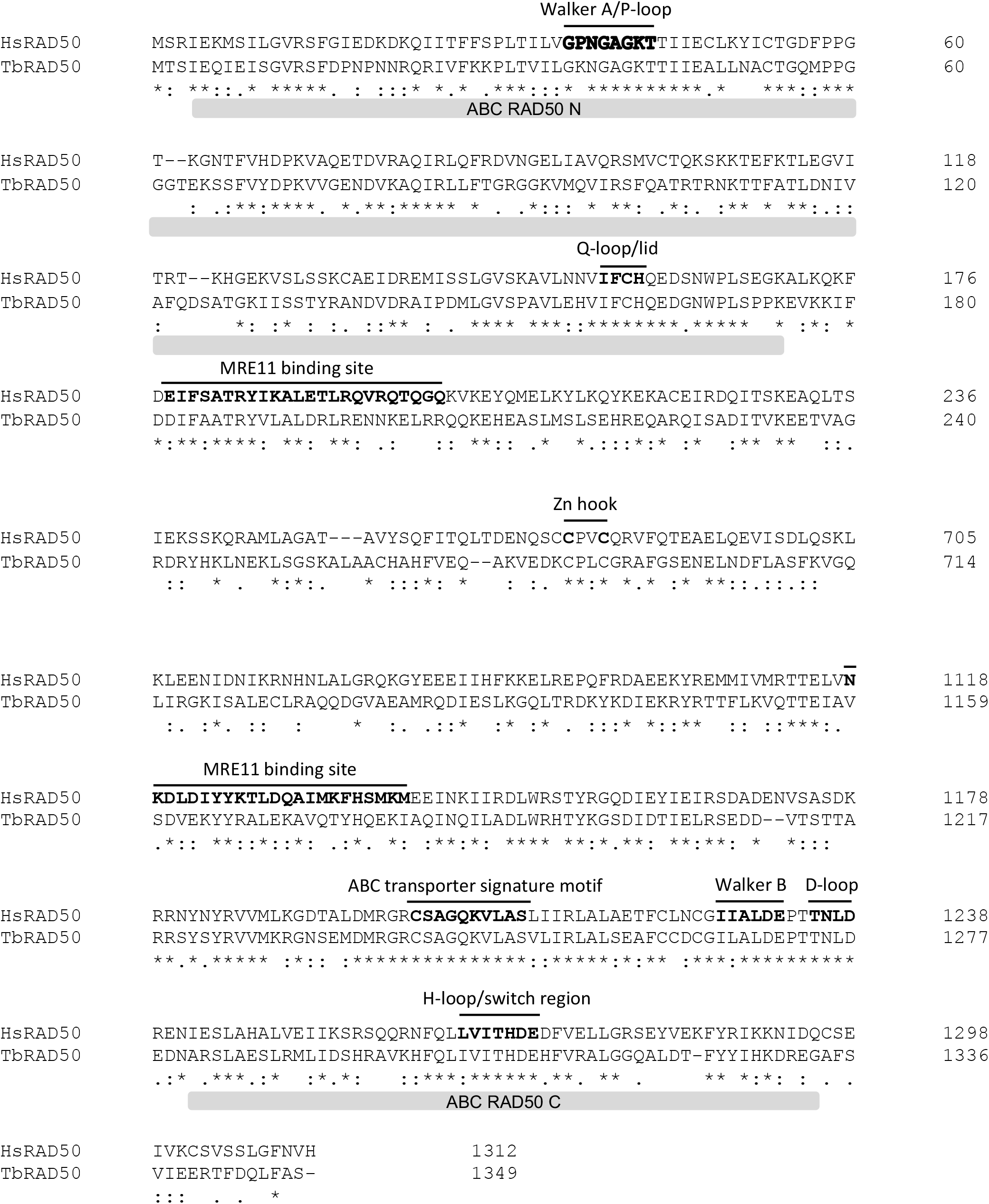
Sequence alignment of the *T. brucei* RAD50 (*Tb*RAD50) with *Homo sapiens* (*Hs*RAD50; UNIPROT: Q92878). Conserved domains are highlighted in bold and the ABC/ATPase domains highlighted with grey bar. Highly conserved residues are highlighted with a * and residues with conserved properties with ‘:’ or ‘.’.

**Supplementary figure 2:**
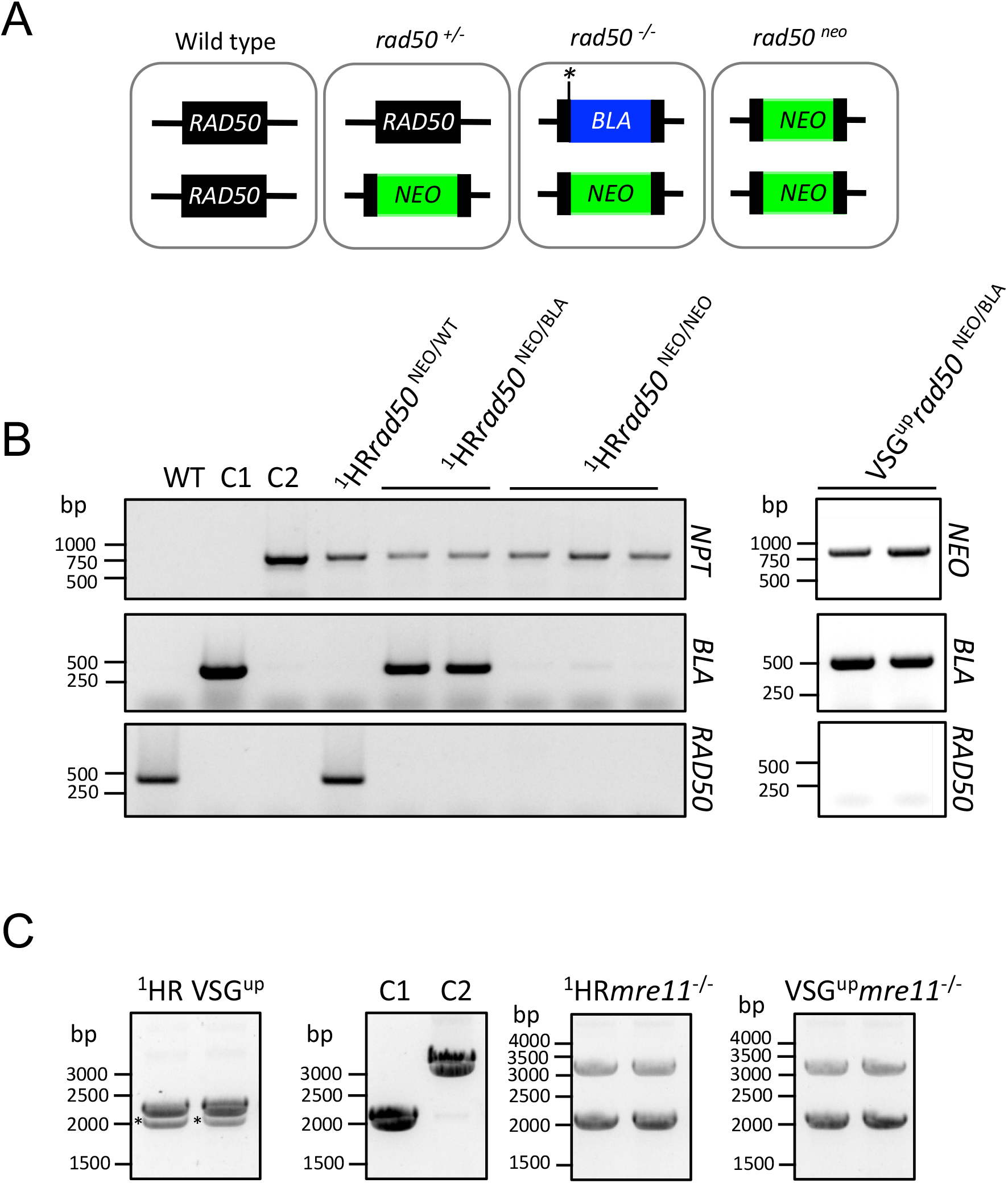
Generation of *rad50* and *mre11* null strains. (A) Upper panel: Schematic showing the knockout strategy to generate the chromosome-internal ^1^HR*rad50* null cell line. (B) PCR assay confirming *RAD50* double allele replacement. (C) PCR assay confirming *MRE11* double allele replacement. * background band. *NEO, Neomycin Phosphotransferase; BLA, Blasticidin deaminase*. C1, control plasmid for *BLA*; C2, control plasmid for *NEO*.

**Supplementary figure 3:**
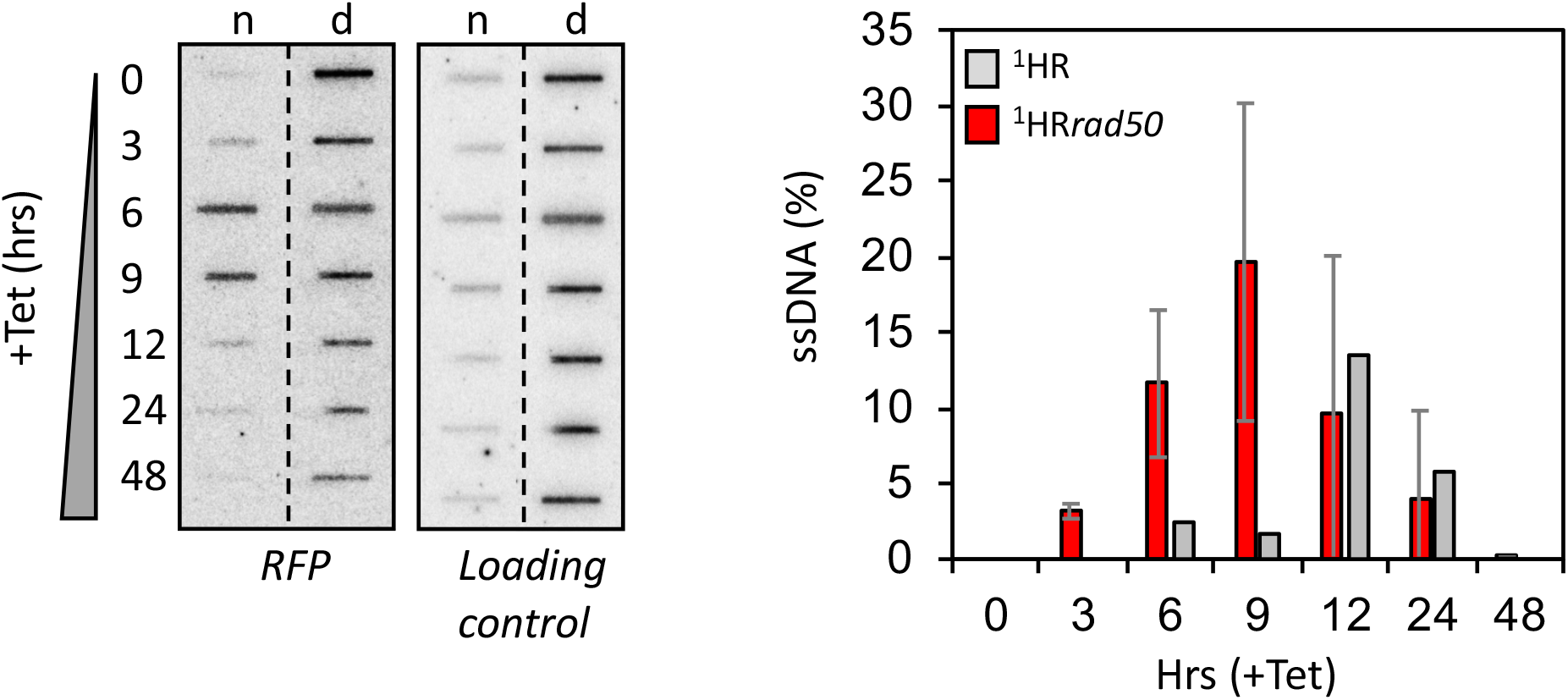
Resection is compromised in the ^1^HR*rad50* nulls Left panel: Biological replicate showing accumulation of ssDNA was monitored using slot-blots, n=2 for biological replicates. Right panel: Quantitative analysis the amount of ssDNA. Error bars, SD for biological replicates for the *rad50* null strains; n=2.

**Supplementary figure 4:**
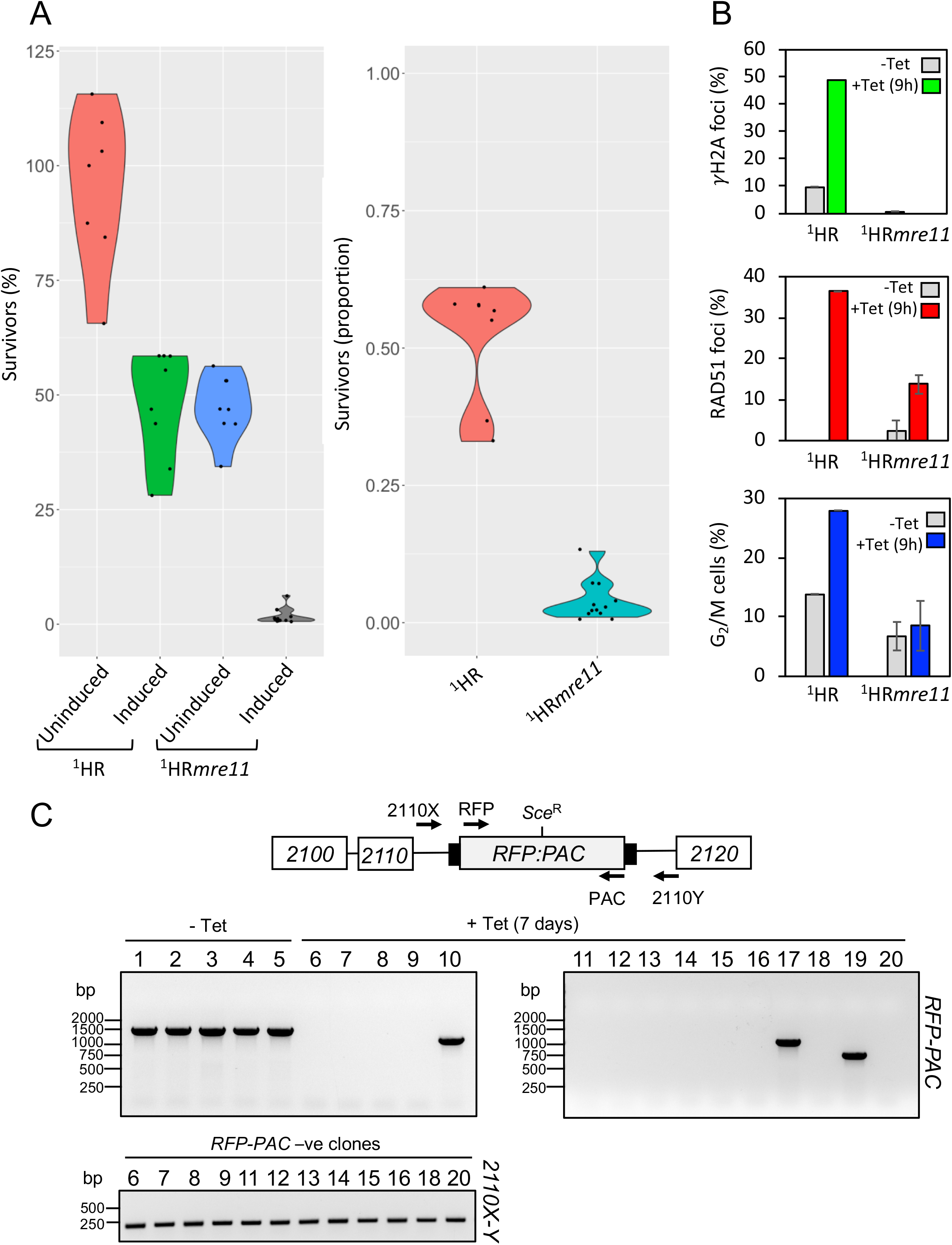
MRE11 is essential for DSB response and repair at a chromosome-internal locus. (A) Clonogenic assay reveals survivors following a DSB at a chromosome-internal locus in the parental and ^1^HR*mre11* null cell lines. Details as in Figure 1 ^1^HR technical replicates; n=2, and with ^1^HR*mre11* biological replicates for the strains; n=2. (B) Upper panel: Immunofluorescence assay to monitoring γH2A foci. The number of positive nuclei were counted in uninduced cells and 12 hours post DSB. Middle panel: Immunofluorescence assay to monitoring RAD51 foci. The number of positive nuclei were counted in uninduced cells and 12 hours post DSB. Lower panel: The number of cells in G2/M phase cells was counted by DAPI staining at several points following induction of an I-*Sce*I break in. G2 cells contain one nucleus and two kinetoplasts. Error bars, SD, for ^1^HR*mre11* biological replicates for the strains; n=2. (C) PCR analysis of repaired subclones. Upper panel: Schematic showing the *2110* locus and position of the *Sce* recognition site (Sce^R^). Position of primers indicated by arrows. Primer sequence detailed in materials and methods. Middle panel: PCR assay of repaired subclones showing *RFP:PAC* presence or absence. Lower panel: PCR assay of repaired subclones that were negative for *RFP:PAC*. Arrows indicate position of primers. White box, genes; Grey box, *RFP:PAC* fusion gene; black box, UTRs.

**Supplementary figure 5:**
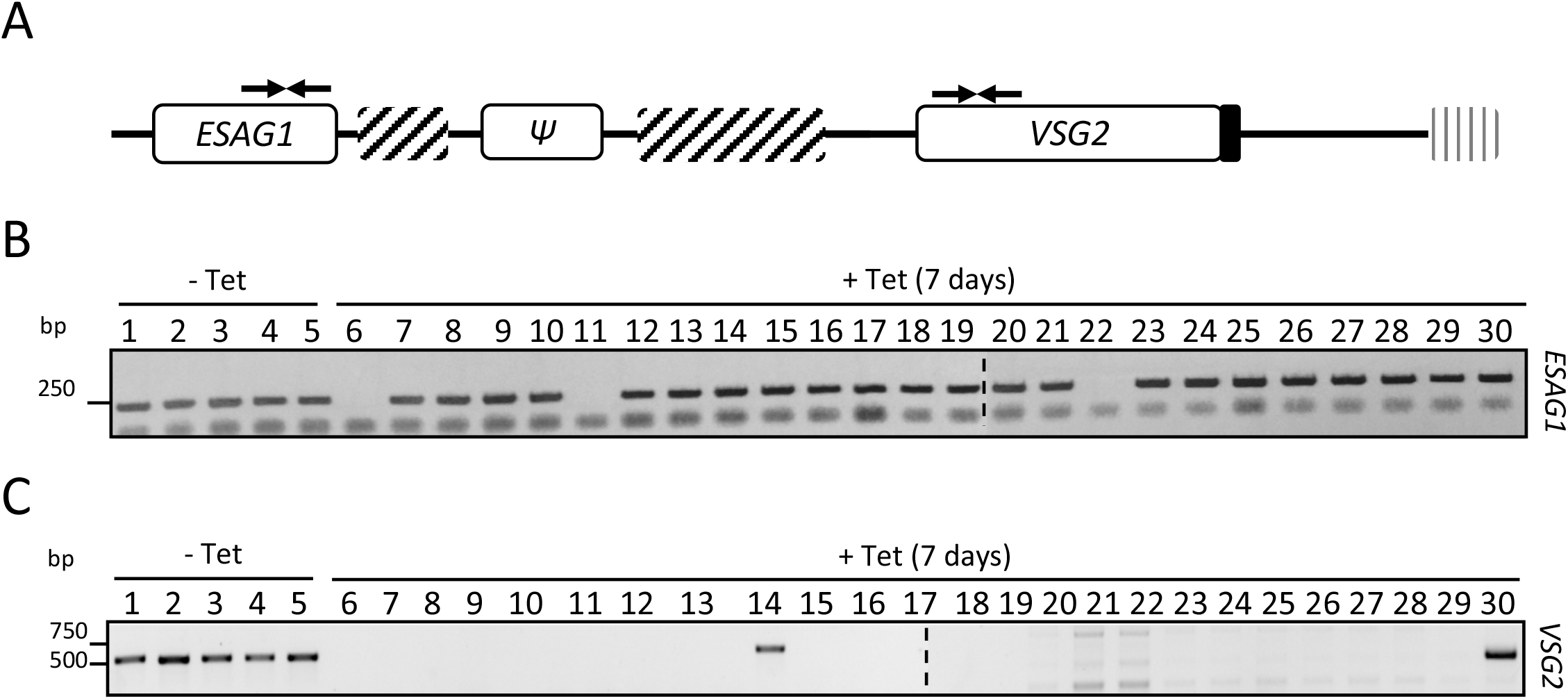
Analysis of VSG^up^*RAD50* strains. (A) Schematic showing the telomeric end of the *VSG2* BES. (B) Presence or absence of *ESAG1* in the repaired subclones. (C) Presence or absence of *VSG2* in the repaired subclones. Arrows indicate position of primers; box with diagonal lines, 70-bp repeats; white boxes, genes; *ψ*, *VSG* pseudo gene; vertical lines, telomere.

**Supplementary figure 6:**
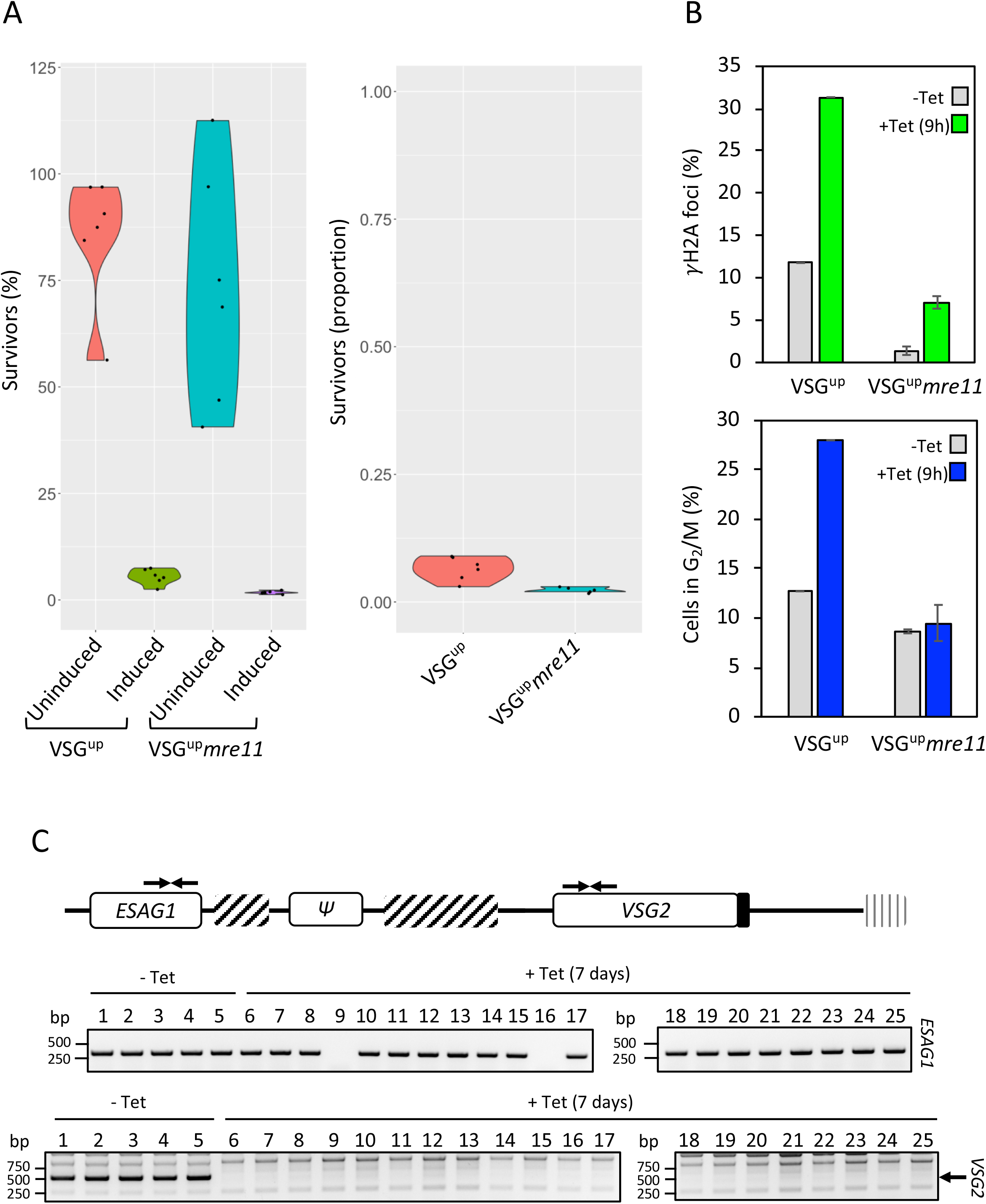
MRE11 is essential for DSB response and repair at an expression site. (A) Clonogenic assay reveals survivors following a DSB in the VSG^up^ strain in the parental and VSG^UP^*mre11* cell lines. Details as in Figure 1 (B) Upper panel: Immunofluorescence assay to monitoring γH2A foci. The number of positive nuclei were counted in uninduced cells and 12 hours post DSB. Lower panel: The number of cells in G2/M phase cells was counted by DAPI staining at several points following induction of an I-*Sce*I break in. G2 cells contain one nucleus and two kinetoplasts. Error bars, SD, for VSG^up^*mre11* biological replicates for the strains; n=2. (C) PCR analysis of repaired subclones. Upper panel: Schematic showing BES1. Position of primers indicated by arrows. Primer sequence detailed in materials and methods. Middle panel: PCR assay of repaired subclones showing *ESAG1* presence or absence. Lower panel: PCR assay of repaired subclones that were negative for VSG2 by immunofluorescence. Arrows indicate position of primers. White box, genes;; ψ, pseudo gene; lined box, 70 bp repeats; black box, UTRs. VSG^up^*rmre11* biological replicates for the strains; n=2.

**Supplementary figure 7:**
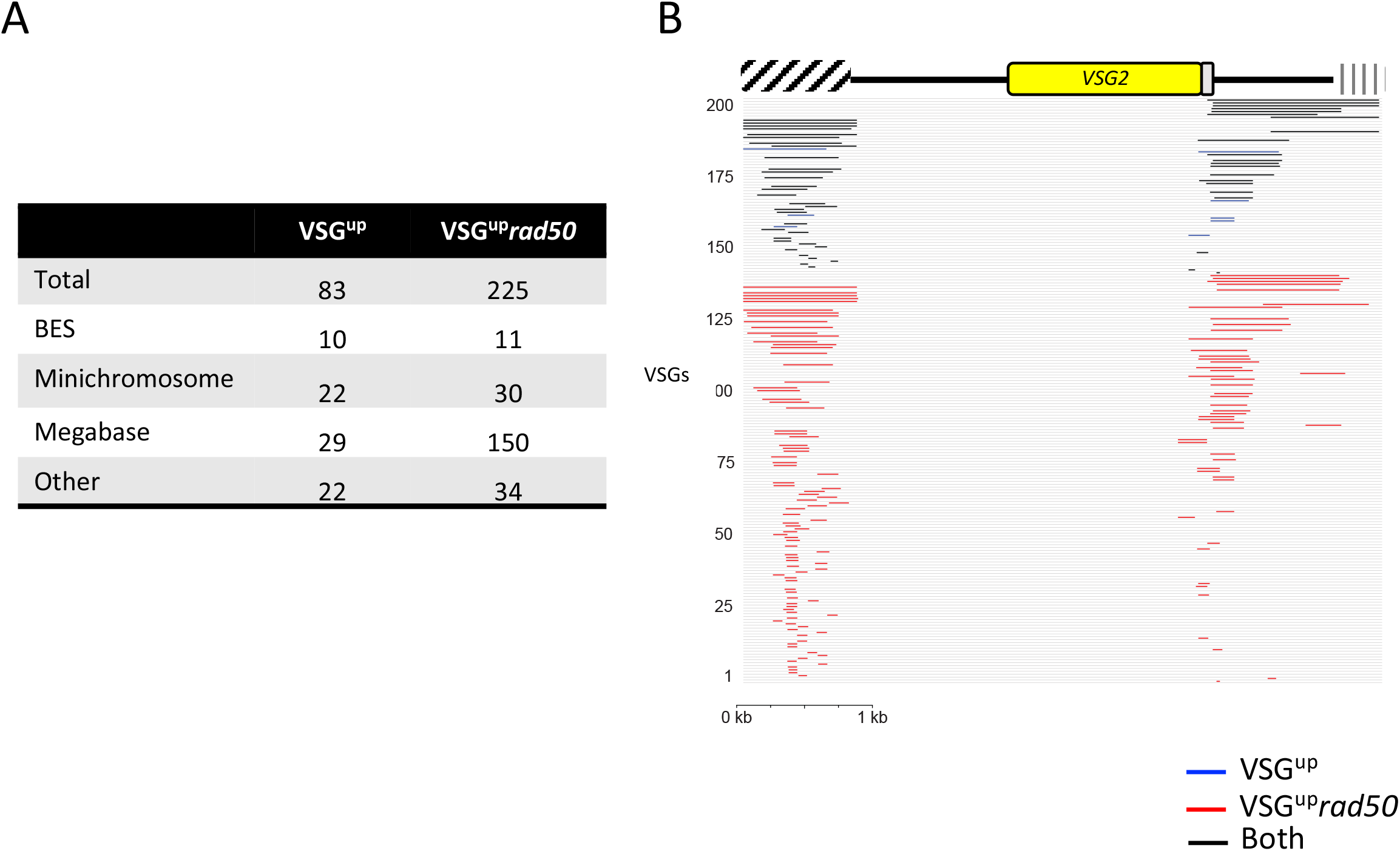
(A) Table showing position of the *VSG* genes used for recombination (C) Proportion of VSG2 in the populations before and after induction of a DSB. (D) Schematic represents the telomeric end of the *VSG2* ES. BLAST analysis of significantly up-regulated genes. BLAST hits are represented as lines showing their position on the *VSG2* query sequence. Black lines – *VSG* genes significantly enriched in both VSG^up^ and VSG^up^*rad50*, blue bars – VSG^up^ only, red bars – VSG^up^*rad50* only.

